# Fibroblast diversification is an embryonic process dependent on muscle contraction

**DOI:** 10.1101/2024.03.18.585498

**Authors:** Lavi Coren, Shelly Zaffryar-Eilot, Anas Odeh, Anna Kaganovsky, Peleg Hasson

## Abstract

Fibroblasts are the most common cell type found in connective tissues, known to play major roles in development, homeostasis, regeneration and disease. Although specific fibroblast subpopulations were associated with different biological processes, the mechanisms and unique activities underlying their diversity has not been thoroughly examined. Turning to skeletal muscle development, we set to dissect the variation of muscle-resident fibroblasts (mrFibroblasts). Our results demonstrate mrFibroblasts diversify following the transition from embryonic to fetal myogenesis prior to birth. We find mrFibroblast segregate into two major subpopulations occupying distinct niches, with interstitial fibroblasts residing between the muscle fibers, and delineating fibroblasts sheathing the muscle mass. We further show these subpopulations entail distinct cellular dynamics and transcriptomes. Notably, we find mrFibroblast subpopulations exert distinct regulatory roles on myoblast proliferation and differentiation. Finally, we demonstrate this diversification depends on muscle contraction. Altogether, these findings establish mrFibroblast diversify in a spatio-temporal embryonic process into distinct cell types, entailing different characteristics and roles.

## INTRODUCTION

Recent advances in single cell technologies have revealed new layers of complexities profiling cell identities, revealing previously thought homogeneous cell populations to be composed of multiple subpopulations with distinct transcriptomes. One such cell type are the fibroblasts, the most common cell type residing in connective tissues, considered as the main source for extracellular matrix (ECM) deposition. Fibroblasts are mostly viewed as undifferentiated cells of mesenchymal origin that can be driven into more defined cell types (e.g., smooth muscle cells, adipocytes or chondrocytes) in response to specific stimuli. They populate all tissues in our bodies, are present and play key roles in virtually all processes from early stages of embryonic development through adulthood to the aging tissue. However, recent observations have demonstrated that fibroblasts are heterogeneous groups of cells with distinct transcriptomes, suggesting that each of these subpopulations has unique roles (Buechler et al., 2021; Plikus et al., 2021; Pradhan et al., 2021; Soliman et al., 2021; Sorrell & Caplan, 2009). Yet how these fibroblasts diversify and what are the specific roles the various subpopulations play, remain largely unknown.

Fibroblasts are identified in the adult by the cell surface markers stem cell antigen-1 (Sca1 or Ly6A/E) and platelet-derived growth factor receptor α (PDGFRα). They are known to have organ-specific functions in determining morphology and differentiation of other cells in the tissue. Fibroblasts, or fibro-adipogenic progenitors (FAPs) are key cells intimately linked with the muscle, essential to all stages of myogenesis from the early embryo through regeneration in the adult, residing in and producing the muscle connective tissue (MCT) which surrounds, protects, and interconnects the primary components of the musculoskeletal system. They are best known for being the main ECM secreting and modifying cells and as such delineate muscle topography by providing essential niches and positional information via biomechanical and biochemical ques (Besse et al., 2020; Hasson, 2011; Hasson et al., 2010; Kardon et al., 2003; Plikus et al., 2021). However, recent studies have demonstrated that fibroblast carry out multiple additional instructive roles including cytokine and growth factor secretion, and satellite cell niche remodeling, which in turn regulates myogenic cell proliferation, progenitors’ self-renewal, differentiation, fusion and patterning (Molina et al., 2021; Nassari et al., 2017; Theret et al., 2021).

Following birth, neonatal myogenesis commences, a phase in which the muscles rapidly grow and mature. By this stage, however, all patterning processes have already terminated (Besse et al., 2020; Biressi et al., 2007; Nassari et al., 2017; Rossi & Messina, 2014). Embryonically, fibroblasts were found to emerge from multiple lineages which converge to form fibroblasts that populate different organs. The limb fibroblasts, which populate the MCT are derived from the lateral plate mesoderm (LPM), where Kardon and colleagues in the early 2000s identified them as a population of TCF7L2/TCF4+ [(transcription factor 4; also known as transcription factor 7 like 2; TCF7L2) expressing cells in the chick and mouse (Kardon et al., 2003; Mathew et al., 2011; Murphy et al., 2011). Other markers expressed by connective tissue progenitors in the mouse embryo include, T-box transcription factors (TBX) 3/4/5 (Colasanto et al., 2016; Hasson et al., 2010), HOX11 (Swinehart et al., 2013) and Odd skipped-related 1 (Osr1) (Stricker et al., 2012; Stumm et al., 2018; Vallecillo-García et al., 2017), which are critical to set up a prepattern that plays an important role for determining myogenic cell differentiation in the limb as deletion of these genes leads to limb muscle embryonic patterning defects (Nassari et al., 2017).

Embryonic development of skeletal muscles is largely divided into two major phases – embryonic (primary) and fetal (secondary) myogenesis. During embryonic myogenesis (in the limb ∼E9.5-E13.5), Pax3 expressing myogenic progenitors migrate from the somite into the nascent limb bud, where they proliferate and commit to a myogenic program, expressing Pax7/Myod1/Myf5 and subsequent myogenic factors to form the muscle preplan. This primary stage in myogenesis is complemented by myoblast and myocyte fusion to form primary myofibers, while individual muscle bundles are formed by MCT orchestrated splitting of the muscle mass. During fetal myogenesis (∼E14.5-P0), fusion and myofiber formation are significantly enhanced, sarcomeres mature, myofibers anchor along the muscle-tendon junctions (MTJs) and specify their twitching type, facilitating the commencement of regulated muscle contraction (Besse et al., 2020; Biressi et al., 2007; Nassari et al., 2017; Rossi & Messina, 2014).

The identification of additional markers for fibroblasts has revolutionized the field and facilitated advancement in our understanding of these key cells as dissecting the specific roles carried out by fibroblasts has been significantly delayed due to the lack of more specific markers. Recent observations demonstrated multiple shared fibroblast subpopulations are present in the various tissues of our bodies, including skeletal muscles (Buechler et al., 2021). In adult muscle, McKellar’s integrative analysis of multiple SC studies have found three FAP subpopulations, however, as many studies report a different number of subpopulations and adopt different nomenclature, consensus in the field has yet to be achieved (Collins & Kardon, 2021; McKellar et al., 2021). Moreover, involvement of specific subpopulations in regulating diverse and muscle specific pathologies was recently noted (Davidson et al., 2021; Henderson et al., 2020; Scripture-Adams et al., 2022; Suárez-Calvet et al., 2023) altogether reinforcing the notion these fibroblast subtypes carry distinct activities in health and disease. These have highlighted the need to explore the underlying heterogeneity of fibroblasts and the roles played by these supposed subpopulations.

Here, we set to explore the diversification of fibroblast subpopulations. We identified fibroblast subpopulations corresponding to Pi16+ and Col15a1+ FAPs (McKellar et al., 2021) to be present in neonate muscles where each one entails distinct, somewhat opposing, myogenic regulatory activities. Focusing on the two largest populations, we find fibroblasts diversify during embryonic development, coinciding with the initiation of muscle contraction following the transition from primary to secondary myogenesis. This process spatially segregates each subtype to a specific niche, failing in paralyzed non-contractile muscles. We suggest the distinct localization of muscle resident fibroblast subpopulations, together with their diverse gene expression and cell dynamics, can facilitate distinct functions and subsequent roles in embryonic muscle development, and possibly in the adult.

## RESULTS

### Muscle fibroblast diversification is an embryonic process

Fibroblast activities are associated with numerous biological processes including embryonic development, homeostasis, tissue regeneration, and aging, but also with multiple pathologies (Buechler et al., 2021; Contreras et al., 2021; Pradhan et al., 2021). The recent advancements in single cell technologies have not only demonstrated the large diversity of fibroblasts (Buechler et al., 2021) but have also highlighted the unexplored distinct activities carried by them in different biological settings. Notably, involvement of specific subpopulations in regulating diverse pathologies that was recently noted (Davidson et al., 2021; Henderson et al., 2020), highlights the need to explore the underlying mechanisms and processes governing this diversity.

To test when fibroblasts diversify into distinct subpopulations, we monitored single cell data of neonate P0 mouse limb muscles (<u>GSE168153</u>) (Yaseen et al., 2021) and focused on the fibroblasts. Among the identified cell populations, five (Fig. 1A) were found to express transcriptional hallmarks of fibroblasts (*Pdgfrα* and *Col1a1*). Interestingly, none of these clusters could be associated with the specific expression of one of the FAP/fibroblasts classical markers (e.g., *Osr1* or *Tcf7l2*) (Fig. 1B) suggesting these markers do not induce a specific subpopulation identity. Each cluster expressed a unique transcriptional signature and will be referred by marker genes: *Smoc2+*, *Ly6a+*, *Gdf10+*, *Cldn1+*, and *Pax7*+/ dual identity fibroblasts (Fig. 1C-E, Fig. S1A) (Yaseen et al., 2021).

**Figure 1.**
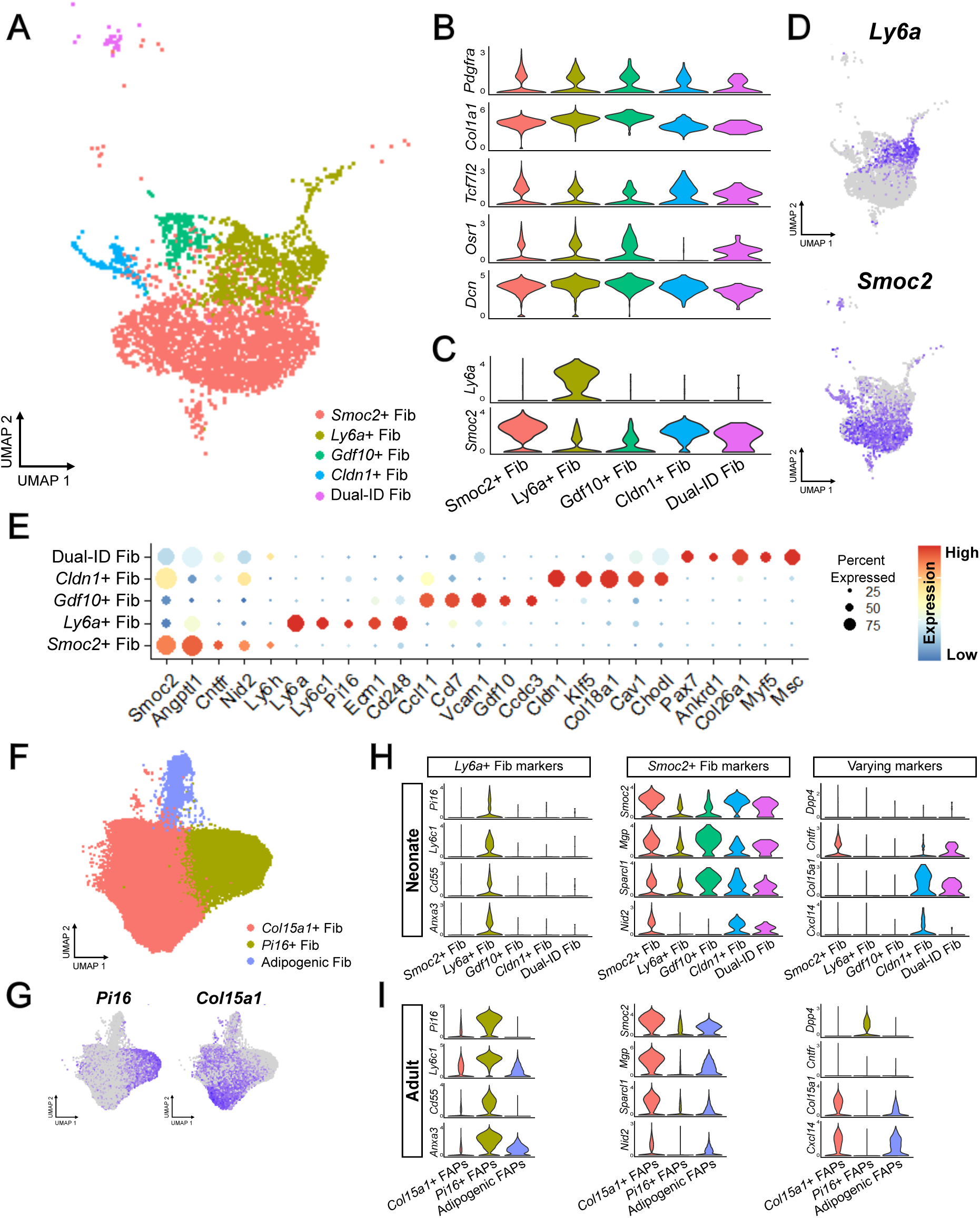
Differential gene expression in muscle fibroblast clusters highlights similarities between neonate and adult subpopulations. (A) UMAP of 5 FAP/fibroblast subpopulations demonstrating discrete clustering at P0. (B) Violin plots for fibroblast hallmark genes demonstrate cells in the 5 clusters are fibrogenic. (C-D) Violin plots (C) and UMAP gene expression (D) for *Ly6a* and *Smoc2* demonstrate specific expression of the two markers in distinct clusters. (E) Dot plot of marker genes of each cluster at P0. (F) UMAP of 3 FAP clusters in adult muscle (data taken from *McKellar et al.,*) (G) Expression of *Pi16* and *Col15a1* demonstrate specific expression of the two markers in distinct adult clusters. (H-I) Violin plots at P0 (H) and adult (I) for differential gene expression (DEG) in 5 FAP/fibroblast clusters at P0 and 3 FAP adult populations showing multiple marker genes’ expression is maintained in the neonate and adult clusters (left and middle columns) while other marker gene expression is changed with age (right column).

Having seen that already at birth fibroblast subpopulations can be observed, we wished to compare these neonate subpopulations to adult muscle resident fibroblasts, where two ’universal fibroblasts’ subpopulations were recently characterized (Buechler et al., 2021) and termed as ’stem’ (known as *Pi16*+\*Dpp4*+) and ’pro-remodeling’ (*Col15a1*+\*Cxcl14*+) fibroblasts (Fig. 1F,G) (McKellar et al., 2021). Remarkably, the two largest fibroblast subpopulations we identify in our analysis, *Smoc2+* and *Ly6a*+, correspond to these ’pro-remodeling’ (expressing *Smoc2*, *Mgp*, *Nid2*, and *Sparcl1*) and ’stem’ (expressing *Pi16*, *Ly6c1*, *Cd55* and *Cd248*) ‘universal fibroblasts’, respectively (Fig. 1H,I and Fig. S1B-D). On the contrary, no expression of key adipogenic markers was found in neonate fibroblasts (Fig. S1E). Although the expression of some markers varies between the neonate (our data) and the adult [e.g., *Ly6a*, *Dpp4*, *Col15a1*, *Cxcl14* (Fig.1H,I)], the results suggest that the main muscle resident FAP/fibroblast subpopulations are largely formed and present already at birth. Altogether, these observations demonstrate fibroblast diversification is an embryonic process.

### Myogenesis is differentially regulated by fibroblast subpopulations

Muscle resident fibroblasts have previously been demonstrated to participate in the regulation of myogenesis in vivo and in vitro (Fiore et al., 2016; Mathew et al., 2011; Murphy et al., 2011). Single cell analyses further suggested fibroblast subpopulations to distinctly interact with myogenic cells during regeneration (McKellar et al., 2021).

To further characterize neonate *Smoc2*+ and *Ly6a*+ fibroblasts, we turned to in-vitro experiments using fluorescently activated cell sorter (FACS) to isolate and culture these subpopulations separately. Taking advantage of the *Prx1Cre* line (Logan & Tabin, 1998) specifically expressed in the lateral plate mesoderm (LPM) derived limb connective tissues, we crossed this Cre line with *ROSA26-nTnG* reporter (Prigge et al., 2013) to label LPM derived cells with GFP nuclei (Yaseen et al., 2021). We sorted Ly6a+ and Smoc2+ fibroblasts (GFP+; CD45-; CD31-) using Sca1 (*Ly6a* encoded protein) and CD200 (highly expressed in Gdf10+ and Cldn1+ fibroblasts) (Fig. 2A and Fig. S2A,B). We collected proportionally fewer Sca1 cells, and verified our sorting using PDGFRα and Sca1 immunostaining (Fig. S2C,D). We further collected myogenic progenitors (MP; satellite cells and myoblasts, tdTomato+; CD45-; CD31-) for culturing (Fig. S2A). An alternative sorting strategy was also applied when sorting fibroblasts isolated from *wild type* (WT) mice pups, using Itga7 (which marks myogenic progenitors) (Fig. S2E,F).

**Figure 2.**
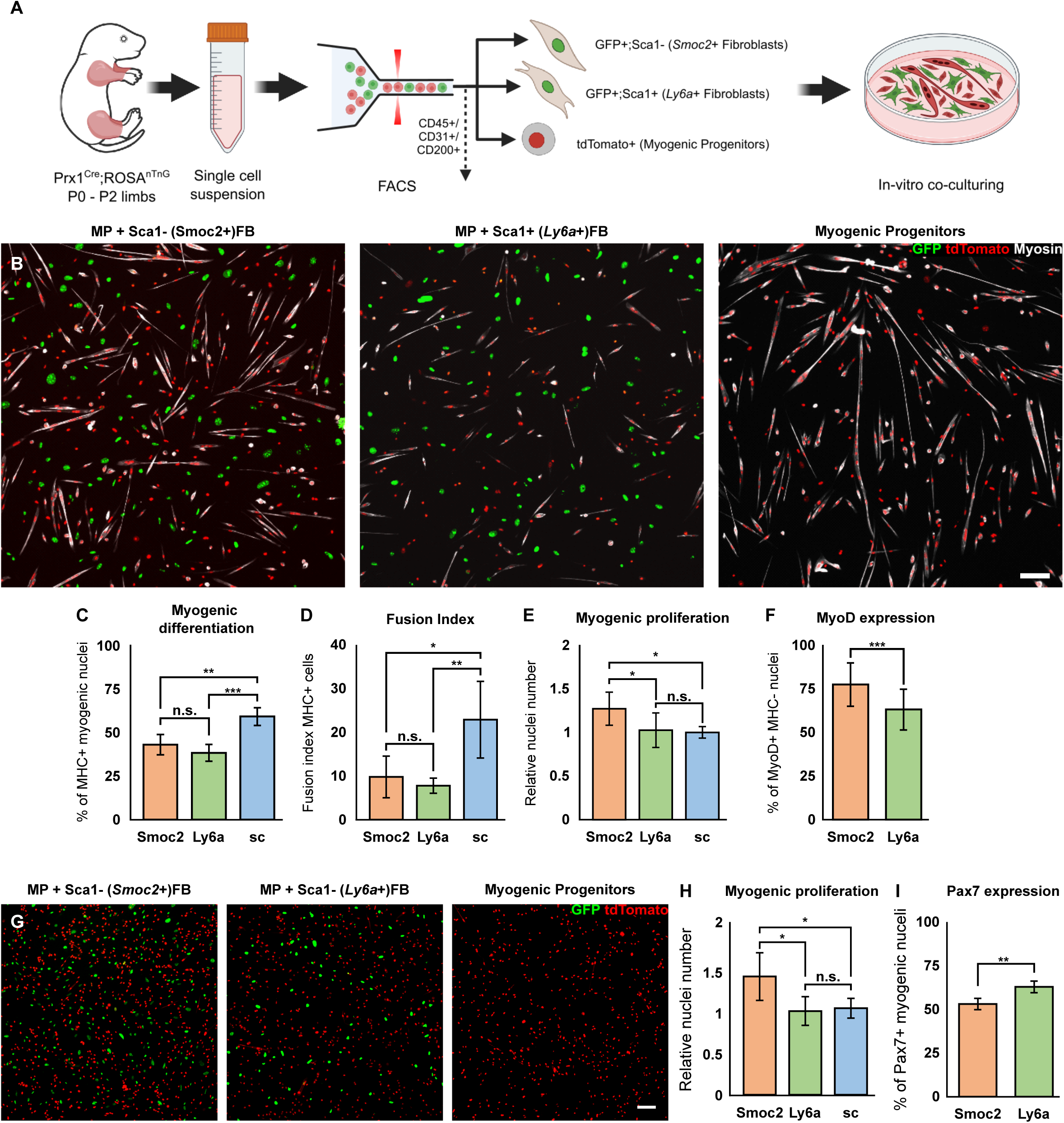
Distinct fibroblasts subpopulations differentially regulate myogenesis in vitro. (A) Schematic of the experimental design for isolation and sorting of fibroblasts subpopulations and myogenic progenitors. (B) Immunofluorescence of myogenic differentiation in-vitro following 72 hours co-culturing of myogenic progenitors with fibroblast subpopulations. Fibroblasts nuceli are GFP+, myogenic nuclei are tdTomato+. GFP (green), tdTomato (red), myosin (gray). Scale bar = µm 100 (C-F) Bar-plot quantification for myogenic differentiation (percentage of MHC+ myogenic nuclei out of all myogenic nuclei) (C), fusion index (D), myogenic proliferation (relative to MP alone in each experiment) (E), and percentage of MyoD expressing undifferentiated (MHC-) myogenic progenitors (F) after 72 hours of differentiation in co-culture. (G) Immunofluorescence of myogenic nuclei abundance in-vitro following 24 hours co-culturing of myogenic progenitors with fibroblast subpopulations. Fibroblasts nuclei are GFP+, myogenic nuclei are tdTomato+. GFP (green), tdTomato (red). Scale bar = µm 100 (H-I) Bar-plot quantification for myogenic differentiation (percentage of MHC+ myogenic nuclei out of all myogenic nuclei) and percentage of Pax7 expressing myogenic progenitors after 24 hours of differentiation in co-culture. (C-E, H-I) Data are presented as mean ± SD. Significance denoted by an asterisk as calculated one-way ANOVA and post hoc Tukey’s test (C-E, H) or two-tailed, unpaired Student’s t test (F,I).

When cultured in-vitro, *Ly6a*+ fibroblasts demonstrated larger cell area and were also significantly more proliferative when compared to *Smoc2*+ fibroblasts in culture (Fig. S2G-I).

Having seen that these two fibroblast subpopulations express distinct transcriptomes and behave differently in vitro, we set to test whether they also functionally regulate myogenesis in a distinct manner. Following sorting, we co-cultured isolated MP with each subpopulation to observe their effect on myogenic differentiation. As control, a similar number of MP were cultured on their own. Following 72 hours of differentiation, we found that when co-cultured with either the *Smoc2*+ or *Ly6a*+ subpopulation, MP underwent significantly less differentiation, evident by less Myosin heavy chain expressing (MHC+) myogenic nuclei and lower fusion index compared to MP cultured on their own (Fig. 2B-D). Remarkably, upon co-culturing MP with the *Smoc2+* subpopulation, significantly more myogenic nuclei were found, suggesting they promoted myogenic proliferation (Fig. 2E). Additionally, an increased proportion of mononuclear myogenic cells co-cultured with *Smoc2*+ fibroblasts expressed MyoD compared to those co-cultured with *Ly6a+* fibroblasts, supporting the idea of diverse effects on myogenesis (Fig 2F). Interestingly, no significant difference was found in the number of fibroblastic nuclei between co-culturing conditions, indicating they proliferated to a similar extent, opposed to their dynamics when cultured alone (Fig. S2J).

To further dissect how fibroblasts affect MP activation, we observed the cells after 24 hours of differentiation. We found significantly more myogenic nuclei when co-cultured with *Smoc2+* fibroblasts, in addition to a lower proportion of Pax7 expressing MP (Fig 2G-I). The observation of these effects after only 24 hours of differentiation suggests they are not exerted solely by distinct ECM deposition, which has yet to sufficiently accumulate. Overall, these results highlight the distinct crosstalk between myogenic cells and each subpopulation, and suggest that *Smoc2*+ fibroblasts promote myogenic proliferation while *Ly6a+* fibroblasts has broad inhibitory effects on MP activation.

### Fibroblasts diversify concomitant with the transition from primary to secondary myogenesis

Embryonic development of skeletal muscles is largely divided into two major phases – embryonic, (primary) [embryonic day (E)9.5-E13.5] and fetal (secondary) [E14.5-P0] myogenesis. During embryonic myogenesis, myogenic progenitors migrate from the somite into the nascent limb bud, proliferate and commit to a myogenic program. This primary stage in myogenesis is complemented by myoblast and myocyte fusion to form primary myofibers, while individual muscle bundles are formed by MCT orchestrated splitting of the muscle mass. During fetal myogenesis, fusion and myofiber formation are significantly enhanced, sarcomeres mature, myofibers anchor along the muscle-tendon junctions (MTJs) and specify their twitching type, facilitating the commencement of regulated muscle contraction.

Our observation that distinct fibroblast subpopulations have already diversified by birth prompted us to question when this process has occurred, and whether it can be attributed to the timing of key developmental myogenic events. Towards this end, we carried out scRNAseq analyses at E12.5 (i.e., towards the end of embryonic myogenesis) and at E15.5 (i.e., at the beginning of fetal myogenesis) of skeletal muscles harvested from both fore and hind limbs of embryos (<u>GSE259340</u>). Following enzymatic digestion and filtration into single cell suspension, cells were sequenced using the 10x genomics platform (Fig. 3A). ScRNAseq data were analyzed using the Seurat R package, while Harmony package was used to integrate the two newly generated datasets with our previously generated P0 dataset (Yaseen et al., 2021) to remove batch effects. Following integration, analysis of the unified dataset has revealed 13 initial clusters, which we annotated based on the expression of canonical cell-type marker genes (Fig. 3B).

**Figure 3.**
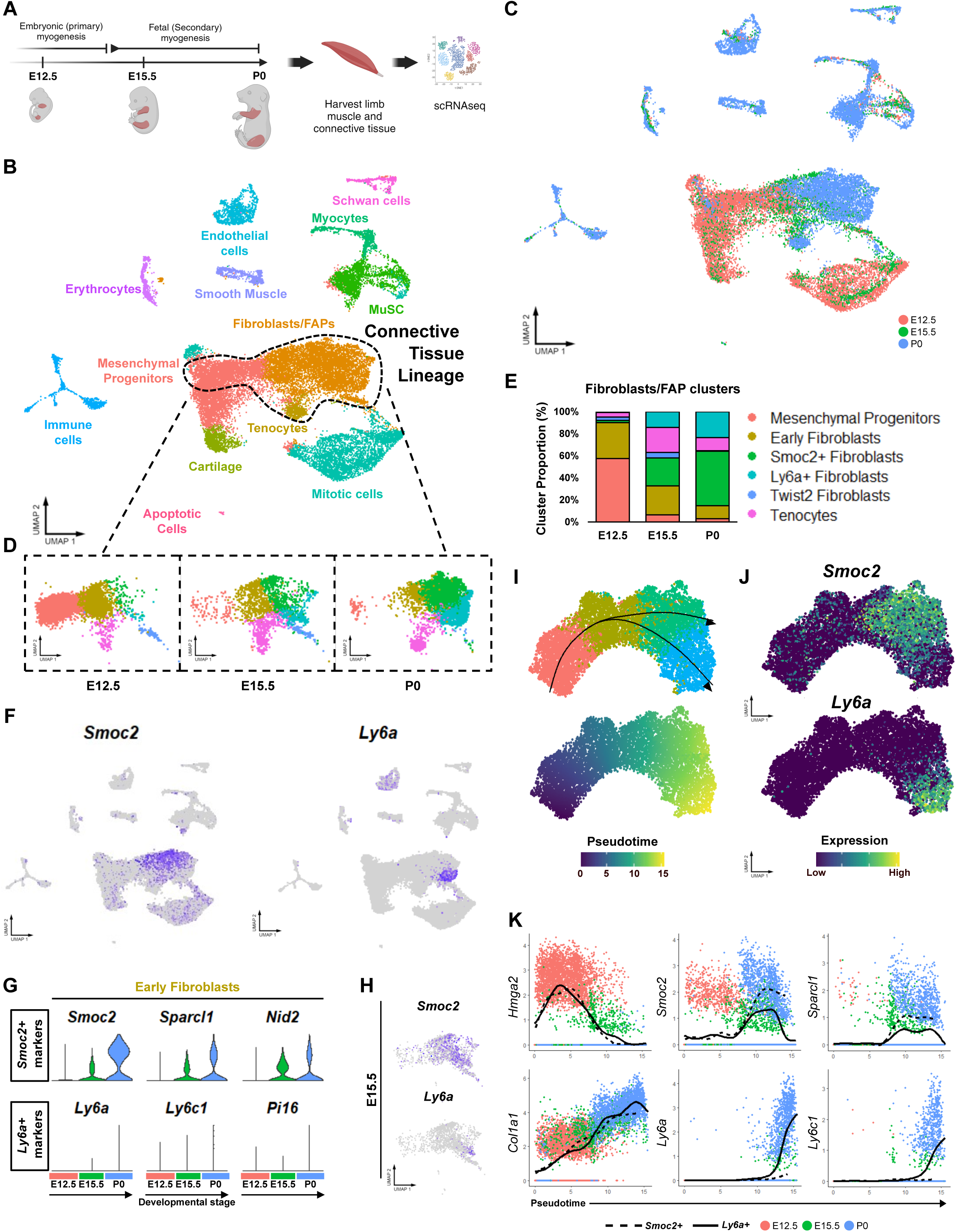
Diversification of FAP/fibroblast is concomitant with the transition from primary to secondary myogenesis. (A) Schematic of the experimental design for scRNAseq of skeletal muscle development. (B) UMAP of three single cell transcriptomal time points (E12.5, E15.5, P0). Fibroblasts/FAP clusters are highlighted with dotted lines. (C) The same UMAP visualization as in (B), showing the time origin of each cell. (D) Connective tissue lineage clusters’ progression at the designated time points. (E) Bar-plot showing changes in the relative proportions of different fibroblasts clusters during development. (F) Expression for *Smoc2+* and *Ly6a+* hallmark markers in integrated dataset. (G) Violin plots for expression of Smoc2+ and Ly6a+ marker genes in ‘Early fibroblasts’ in each timepoint. Colors represent developmental stage of origin as in (C). (H) UMAP showing expression for *Smoc2* and *Ly6a* in E15.5 fibroblasts. (I) Slingshot trajectory and pseudotime visualization of fibroblast diversification across development. Colors represent cluster identity as in (D) or pseudotime score. (J) UMAPs depicting the expression patterns of *Smoc2* and *Ly6a* in fibroblast lineage trajectories during development. (K) Pseudotime ordered single-cell expression trajectories for genes enriched in early (*Hmga2*) and late (*<u>Col1a1</u>*) connective tissue development (left column), in *Smoc2*+ fibroblasts (*Smoc2*, *Sparcl1*), and in *Ly6a*+ fibroblasts (*Ly6a*, *Ly6c1*). Overlaid lines correspond to inferred cell trajectories associated with ending in the *Smoc2*+ (hatched) and *Ly6a*+ (solid) clusters. Colors represent developmental stage of origin as in C.

Fibroblasts were marked based on their expression of *Pdgfra, Col1a1* and *Lumican (Lum)*, common fibroblast markers (Fig. 3B and Fig. S3A,B). Two other connective tissue related populations were detected: tenocytes, marked by the expression of the tenogenic *Scx* transcription factor, and a cluster expressing both fibroblasts and cartilage marker genes that was detected mainly at E12.5 corresponding to previously described ‘Mesenchymal Progenitors’ (Fig. 3B,C and Fig. S2A) (Markman et al., 2023; Reinhardt et al., 2019). Other annotated cell types were two myogenic populations, Muscle Stem Cells (MuSC; expressing *Pax7*) and the more mature myocytes (expressing myogenic transcription factors *Myod1* and *Myog*), along with mitotic proliferating cells, smooth-muscle cells, endothelial, schwan and immune cells (Fig. 3B and Fig. S2A). This dataset enabled us to examine cell populations and gene expression dynamics of fibroblasts during embryonic muscle development.

Next, to analyze the dynamics and differentiation of fibroblasts during embryogenesis, we clustered the connective tissue lineage populations (Mesenchymal Progenitors, Fibroblasts, and Tenocytes) at a higher resolution which revealed 4 fibroblast subpopulations; ’Early Fibroblasts’, *Smoc2*+ fibroblasts, *Ly6a*+ fibroblasts and *Irx5*+ fibroblasts (Fig. 3D,E and Fig. S3C). *Ly6a+* and *Smoc2+* fibroblasts populations corresponded to our previously described P0 ‘universal fibroblasts’ (Fig. 1) (McKellar et al., 2021), expressing their marker genes *Ly6a, Ly6c1* and *Pi16* or *Smoc2, Sparcl1,* and *Cntfr*, respectively (Fig. 3F, Fig. S3C,D). ’Early fibroblasts’ were abundant mainly at E12.5 and E15.5, expressing general fibroblastic markers. A small fibroblast cluster observed mostly at E15.5 highlighted by *Irx5* was further observed (Fig. 3D,E, Fig. S3C). Out of the three minor subpopulations found at P0 (Yaseen et al., 2021, Fig. 1) *Cldn1*+ and *Gdf10*+ [shown by *Ccdc3* expression] fibroblasts clustered together with *Smoc2+* and *Ly6a+* fibroblasts, respectively (Fig. 1E), whereas the ‘dual-identity’ fibroblasts clustered together with the myogenic cells (shown by myogenic cells expressing fibroblastic markers) (Fig. S3B red arrow and Fig. S3E, Cf. Fig. 3B).

To investigate the emergence and diversification of fibroblast subpopulations, we examined the composition of the connective tissue lineage in our data throughout development. At E12.5, the largest fibroblast subpopulation detected were of the ’early fibroblasts’ cluster (32%) which decreased at E15.5 (26%) and P0 (11%), whereas *Smoc2+* and *Ly6a+* fibroblasts emerged at E15.5 (25% and 14% respectively) later expanding at P0 (50% and 23% respectively) (Fig. 3E). Notably, early fibroblasts expressed some *Smoc2+* markers (e.g., *Smoc2*, Sparcl1 and *Nid2*) and showed similar genetic signature to *Smoc2*+ fibroblasts at later stages (Fig. 3G and Fig. S3F), prompting us to view them as *’Smoc2+* like’ fibroblasts. Thus, our data indicates that by E12.5 *’Smoc2+* like’ early fibroblasts were readily found, although they do not express all the hallmark genes of this subpopulation. By E15.5 the majority of *Smoc2+* fibroblasts assume a more mature identity, similar in gene expression to that observed at later stages, where they constituted the majority of fibroblasts (Fig 3E,H-I and Fig. S3F-G). In contrast, *Ly6a*+ fibroblasts were not detected in early embryonic myogenesis, however expression of their marker genes emerged by E15.5, co-expressed with *Smoc2+* and other markers, becoming more defined and distinct by P0 (Fig. 3E,H and Fig. S3F-G). Pseudo-time trajectory analysis of the fibroblast lineage further showed expression of *Smoc2*+ markers in *Ly6a*+ lineage, which diminished following the emergence of *Ly6a*+ marker expression (Fig. 3I-K). These results suggest that during primary myogenesis, the muscle landscape is dominated by ’Smoc2+ like’ fibroblasts, which diversify and split to more defined adult-like *Smoc2+* and *Ly6a+* fibroblasts upon the transition from embryonic to fetal myogenesis.

Notably, expression of top marker genes for the fibroblast subpopulations identified in chick embryos (e.g., *Gap43*, *Mfap5*, and the Annexin family; *Fibin* and *Egfl6*, (Hirsinger et al., co-submitted) and human [*Cd55*, *Mfap5*, *Fbn1*, and *Pcolce2; Smoc2, Sparcl1, Adh1* (Fig. 1E,H and Fig. S3H,I)] (De Micheli et al., 2020a; Fitzgerald et al., 2023) suggest these identities to be evolutionary conserved cell types, indicating they may hold central functional roles in these fibroblast subpopulations.

### Fibroblasts subpopulations reside in distinct niches within the muscle

Our bioinformatics analysis demonstrated that fibroblast diversification is temporally regulated during embryonic development. To test this hypothesis and view this in vivo, we used fluorescent in situ hybridization (FISH) and immunofluorescence analyses at E15.5, the onset of *Ly6a*+ fibroblasts emergence (when fibroblast population is dominated by *Smoc2*+ fibroblasts), and P0 (when *Ly6a+* fibroblasts are more readily observed) (Fig. 3). Strikingly, we found that fibroblast diversification is not only a temporally-but also a spatially-defined process. While at birth, *Pi16+/Ly6a+/*Ly6c+ fibroblasts are delineating the muscles, specifically located in the muscle periphery (epimysium/perimysium) (Fig. 4A, Fig S4A-C), *Smoc2+/Sparcl1+* fibroblasts are hardly expressed in this region and are robustly found in the interstitium adjacent to the myogenic progenitors and in between muscle fibers (Fig. 4A, Fig. S4D). We find this spatial distribution to be maintained also around the myofiber tips, where *Pi16*+ fibroblasts seem to form a continuous sheath over the MTJ (Fig. 4B,C). Interstingly, by E15.5 not only are *Pi16+* cells harder to find, whereas *Smoc2+* cells are found in the muscle periphery, a region by which at P0 is solely occupied by *Pi16*+ cells (Fig. 4D). These observations prompted us to ask whether the transition of peripheral fibroblast identity occurs via ‘Identity transition’ i.e., direct change of cell identity of all peripheral fibroblasts, or whether it arises via a ’replacement’ mechanism where a few *Smoc2*+ “progenitors” switch on a *Ly6a*+ program, and these then proliferate and populate the muscle periphery. To address this question, we looked at E16.5, an intermediate stage in fibroblast diversification; we found an abundance of double-positive (*Smoc2*+ and *Pi16*+) cells delineating the muscles (Fig. 4E,E’). This concurrent expression of *Smoc2* and *Ly6a* are in accord with our expression-over-pseudotime results (Fig. 3K), support the notion of identity transition for achieving fibroblast heterogeneity. Remarkably, when we examined the expression of *Smoc2* at an even earlier stage (E13.5) when ’*Smoc2+* like’ early fibroblasts are abundant (Fig. 3), we found the majority of fibroblasts to be found lining the muscle mass, where *Smoc2* expression was also detected (Fig. 4F,G). These results correspond to findings in avian development (Hirsinger et al., co-submitted), suggesting that both fibroblast subpopulation identities as well as their spatiotemporal diversification during embryonic development are evolutionary conserved events.

**Figure 4.**
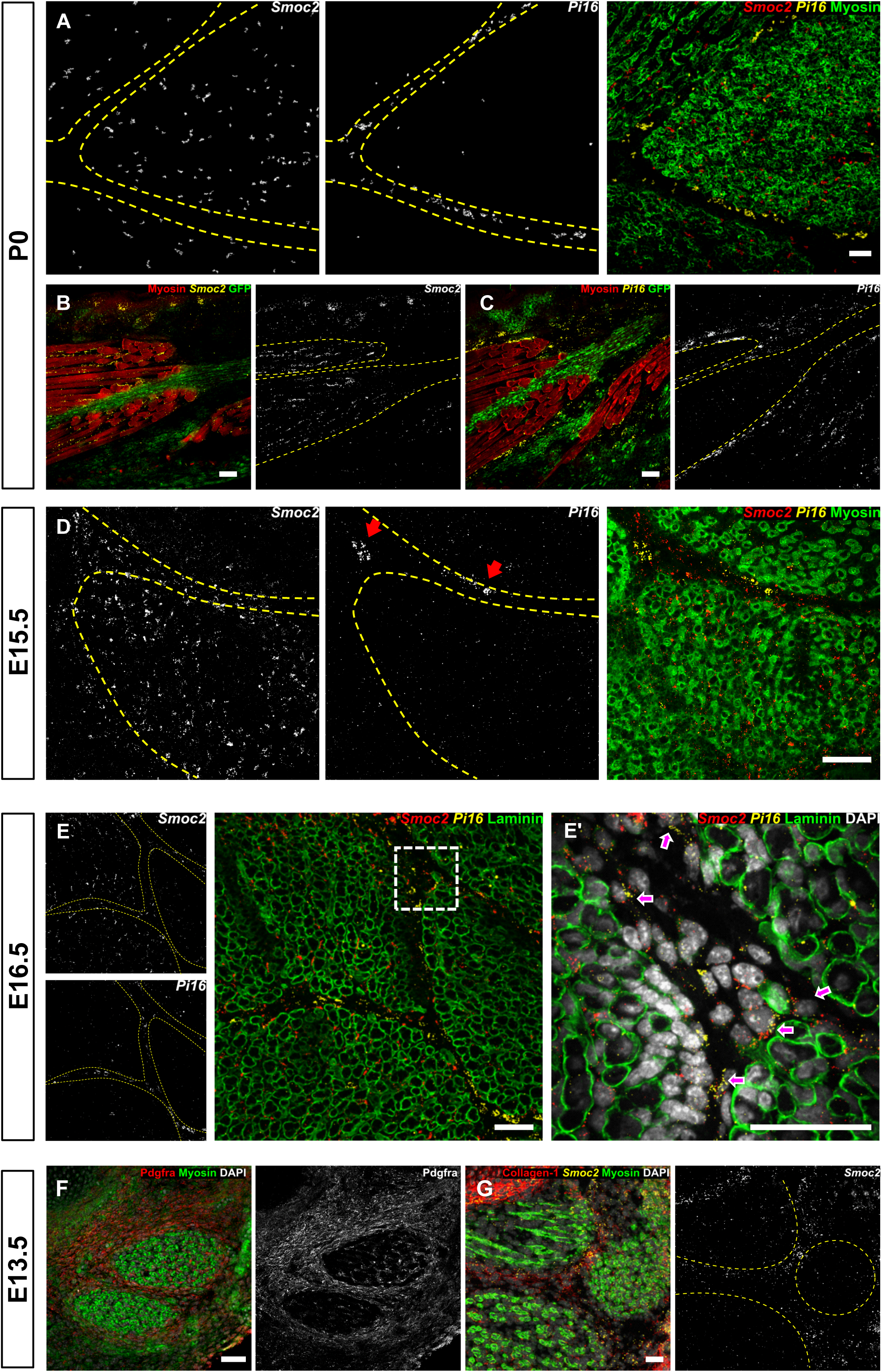
*Smoc2+* and *Pi16+* fibroblasts distinct expression patterns shift during development. (A-G) Immunofluorescence staining and RNA FISH of fibroblast subpopulation markers (*Smoc2* and *Pi16*) in muscle cross sections across development. The muscle borders are highlighted by an orange dashed line. Scale bars = 50 µm (A) P0 muscle cross section. Myosin (green) *Smoc2* (red) *Pi16* (yellow). (B-C) *Prx1^Cre^; ROSA^nTnG^*P0 sagittal sections of the MTJ region. GFP (green) Myosin (red) and *Smoc2* (B) or (*Pi16*) (C) (yellow). (D-E’) E15.5 (D) and E16.5 (E-E’) muscle cross section. Myosin (D) or Laminin (E-E’) (green) *Smoc2* (red) *Pi16* (yellow). Arrows highlight *Pi16* and *Smoc2* dual expressing cells. (F-G) E13.5 muscle cross section. Myosin (green), *Smoc2* (yellow) and Pdgfra (F) or Collagen 1 (G) (red).

The interstitial and peripheral niches correspond to distinct MCT layers, the endomysium and perimysium/epimysium, respectively, both of which are highly enriched in ECM. The distinct spatial expression of fibroblast subpopulation raised the hypothesis this distribution may contribute to the unique properties of each MCT layer. To further analyze whether subpopulation specific ECM deposition may drive different characteristics of the MCT layers, we compared the expression of ECM and ECM related genes (Shao et al., 2020, 2023) at P0 across the fibroblast subtypes. Based on our transcriptomal data *Ly6a* + fibroblasts were found to express an array of ECM regulators (Fig. S4A,B), including Lysyl oxidase (*Lox*), and ECM glycoproteins (Fig. S4C) such as Elastin (*Eln*), Fibrilins (e.g., *Fbn1*), Fibulins (*Efemp1/Fbln3*) and Microfibril-associated proteins (e.g., *Mfap5*), which are associated with formation of elastic fibers (Kielty et al., 2002). Genes related to hyaluronic acid biosynthesis (*Ugdh*, *Has1*) and modifications (*Tnfaip6*) were also found to be expressed by *Ly6a+* fibroblasts (Fig. S4D). Comparing expression of different functional groups of collagens (Karsdal et al., 2020), revealed *Ly6a*+ fibroblasts express fibrillar collagens (e.g., ColI/XIV), whereas *Smoc2*+ fibroblasts primarily express network-forming collagens (e.g., Col IV/VI subtypes) (Fig. S4E). In addition to collagen IV, *Smoc2+* fibroblasts show expression of nidogen-2 (*Nid2*), both of which are major components of basement membranes where these cells reside (W. Zhang et al., 2021). By imposing our spatial findings on transcriptomal data, these analyses highlight further functional diversity between *Ly6a+* and *Smoc2+* fibroblasts, and provide new insight into the characteristics and composition of different MCT layers.

Out of the five fibroblasts subpopulations we identified at P0 (Fig. 1), three were shown to be spatially associated with distinct niches within the muscle (*Ly6a*+ and *Smoc2*+, this paper; Dual-identity fibroblasts, Yaseen et al., 2021), therefore we speculated whether the two remaining ‘minor’ clusters could also be associated to a specific niche. GO analysis of the top 100 differentially expressed genes in *Gdf10+* fibroblasts (Fig. S6A) have highlighted several terms associated with development of the vascular system (Fig. S6B). Notably, Gdf10+ fibroblasts were found to express the pericyte-specific markers *Ndufa4l2*, *Rgs5* and *Kcnj8* (Fig. S6C) (Arostegui et al., 2022). In line with this analysis, staining of neonate muscles have identified Pdgfra+/*Gdf10+* cells sheathing blood vessels (Fig. S6D-F’), suggesting *Gdf10+* fibroblasts to be ‘pericyte-like’ cells that may have a role in muscle vasculature embryonic development.

Lastly, *Cldn1+* fibroblasts corresponded to perineural fibroblasts described previously in both mice and humans, expressing *Ngfr* and *Fzd2* (Arostegui et al., 2022; Kedlian et al., n.d.; Leinroth et al., 2022; Muhl et al., 2020) (Fig. S6 G,H). Immunostaining for Cldn1 have revealed Cldn1+ fibroblasts to delineate neurons (Milner & Cameron, 2013) in neonate muscles (Fig. S6 I-J). Notably, Schwan cells’ derived *Desert hedgehog* (*Dhh*) has been reported to regulate the structural and functional integrity of peripheral nerves (Parmantier et al., 1999; Sharghi-Namini et al., 2006; Zotter et al., 2022). Accordingly, Cldn1+ fibroblasts express the hedgehog pathway target genes *Ptch1* and *Gli1* (Fig. S6 G,H). Altogether, these results suggest perineural and pericyte-like fibroblasts diversify and become spatially distinct during embryonic development similarly to *Ly6a*+ and *Smoc2*+ fibroblasts.

### Fibroblast subpopulations entail distinct cellular dynamics

Cell division, migration, and inductive signaling are some of the mechanisms known for achieving spatial distribution and patterns of cells during embryonic development (Salazar-Ciudad et al., 2003). At birth, *Smoc2*+ and *Ly6a*+ fibroblasts show a distinct spatial separation; our results suggest this organization is achieved by cell ‘identity transition’ of muscle delineating cells (Fig 4). In order to get a better understanding of the mechanisms involved in this spatiotemporal diversification process, we aimed to look at the dynamics of cell proliferation and how it may contribute to the formation of fibroblast subtypes’ distinct patterns. Immunostaining for the cell cycle marker phospho-Histone H3 (pHH3) demonstrates that by E18.5, *Smoc2*+ fibroblasts are almost three times more proliferative than *Pi16*+ fibroblasts (Fig. 5A-B). To observe these cellular dynamics during development, we compared relative proliferation between interstitial and peripheral cells, as by E15.5 *Pi16*+ cells have yet to differentiate and are scarce. At this stage, i.e. when all fibroblasts are presumably less diverse, we find no significant difference between interstitial and peripheral proliferation (Fig. 5C-E). Our results demonstrate that interstitial proliferation remains at relatively the same level as the embryo develops, while peripheral proliferation decreases significantly as *Ly6a*+ fibroblasts assume their mature identity. These results are further supported by an abundant expression of *Smoc2*+ markers (*Smoc2*, *Angptl1*, *Nid2*) but not those of *Ly6a*+ (*Ly6a*, *Pi16*, *Ly6c1*) in mitotic clusters (Fig. 5F,G). Overall these findings support the model of ‘identity transition’ underlying the conversion of delineating fibroblast identity, and define cell proliferation as an additional distinct characteristic of fibroblasts subtypes.

**Figure 5.**
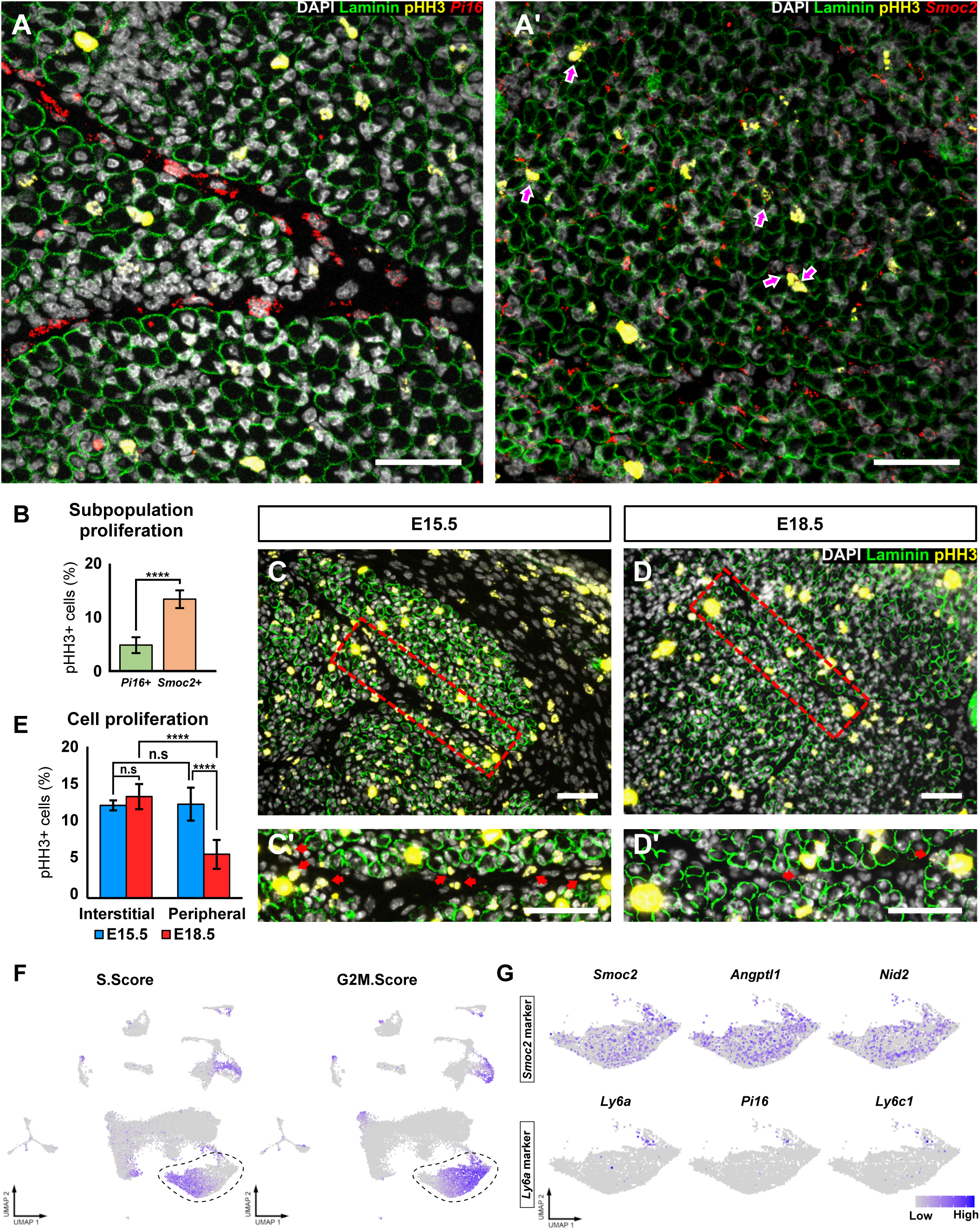
Proliferation of Ly6a+ fibroblasts decreases with development. (A-A’) RNA FISH of fibroblast subpopulation markers (*Smoc2* and *Pi16*) and Immunofluorescence staining for cell cycle marker pHH3 in E18.5 muscle cross sections. DAPI (gray), Laminin (green), pHH3 (yellow) and *Pi16* (A) or *Smoc2* (A’) (red). Arrows highlight pHH3+ fibroblasts. Scale bars = 50 µm (B) Bar-plot quantification for pHH3+ *Pi16*+ or *Smoc2*+ fibroblasts at E18.5 (A-A’). n≥4 per localizations and stage group. Data are presented as mean ± SD. Significance denoted by an asterisk as calculated by two-tailed, unpaired Student’s t test. (C-E) Immunofluorescence staining for cell cycle marker pHH3 (C-D’) in muscle peripheral and interstitial cells at E15.5 (C-C’) and E18.5 (D-D’) muscle cross sections. DAPI (gray), Laminin (green), pHH3 (yellow). Arrows highlight pHH3+ fibroblasts in muscle peripheral cells (C’,D’). Scale bars = 50 µm (E) Quantification for percent of pHH3+ muscle peripheral and interstitial cells at E15.5 and E18.5. n≥4 per localizations and stage group. Data are presented as mean ± SD. Significance denoted by an asterisk as calculated by two-tailed, unpaired Student’s t test. (F) Cell-cycle scoring for the integrated dataset. Mitotic cells cluster highlighted by a dashed line. (G) Expression of characteristic markers for Smoc2+ (upper) and Pi16+ (lower) fibroblasts in mitotic cells cluster in the integrated data.

### Muscle contraction is essential for fibroblast diversification

During muscle development, several key events and processes take place in the embryonic limb. We therefore hypothesized whether one of them could drive fibroblast diversification. Concomitant with the emergence of *Ly6a*+ fibroblasts is the transition from embryonic to fetal myogenesis at around E14.5. Remarkably, this transition is accompanied by maturation of the muscle contractile units, MTJ formation, and initiation of myofiber contractions, which are known to be crucial for limb patterning and muscle transcriptional changes during development (Dos Santos et al., 2023; Felsenthal & Zelzer, 2017; Huang et al., 2015; Lipp et al., 2023). Therefore, to test whether muscle contraction could impact fibroblasts diversification we analyzed limbs of *muscle dysgenic* (*mdg*) embryos, harboring a lethal point mutation in the skeletal muscle specific *Cacna1s* gene which encodes the alpha subunit of Cav1.1 calcium channel, a lethal mutation resulting in paralyzed non-contractile skeletal muscles.

At E18.5, *mdg* embryos are hunchback and smaller than their littermates (Filipova et al., 2018). Whole mount and cross-section staining of the limbs show muscle hypoplasia, albeit all major muscle groups being well-defined and overall muscle patterning is normal (Fig. S7A-B’’). Taking advantage of these mice, we monitored fibroblast subpopulations. Strikingly, we find that muscle peripheral fibroblasts which normally express *Ly6a+* markers (*Pi16*, Ly6a, and Ly6c) no longer express them. Instead, we find these peripheral fibroblasts to express *Smoc2* (Fig. 6A-B’ and Fig. S7C-E’, Cf. Fig. S4). This disruption to the fibroblasts cellular identity raised the question of whether peripheral fibroblasts failed to initiate diversification or whether they lost their cellular identity in lack of muscle contraction.

**Figure 6.**
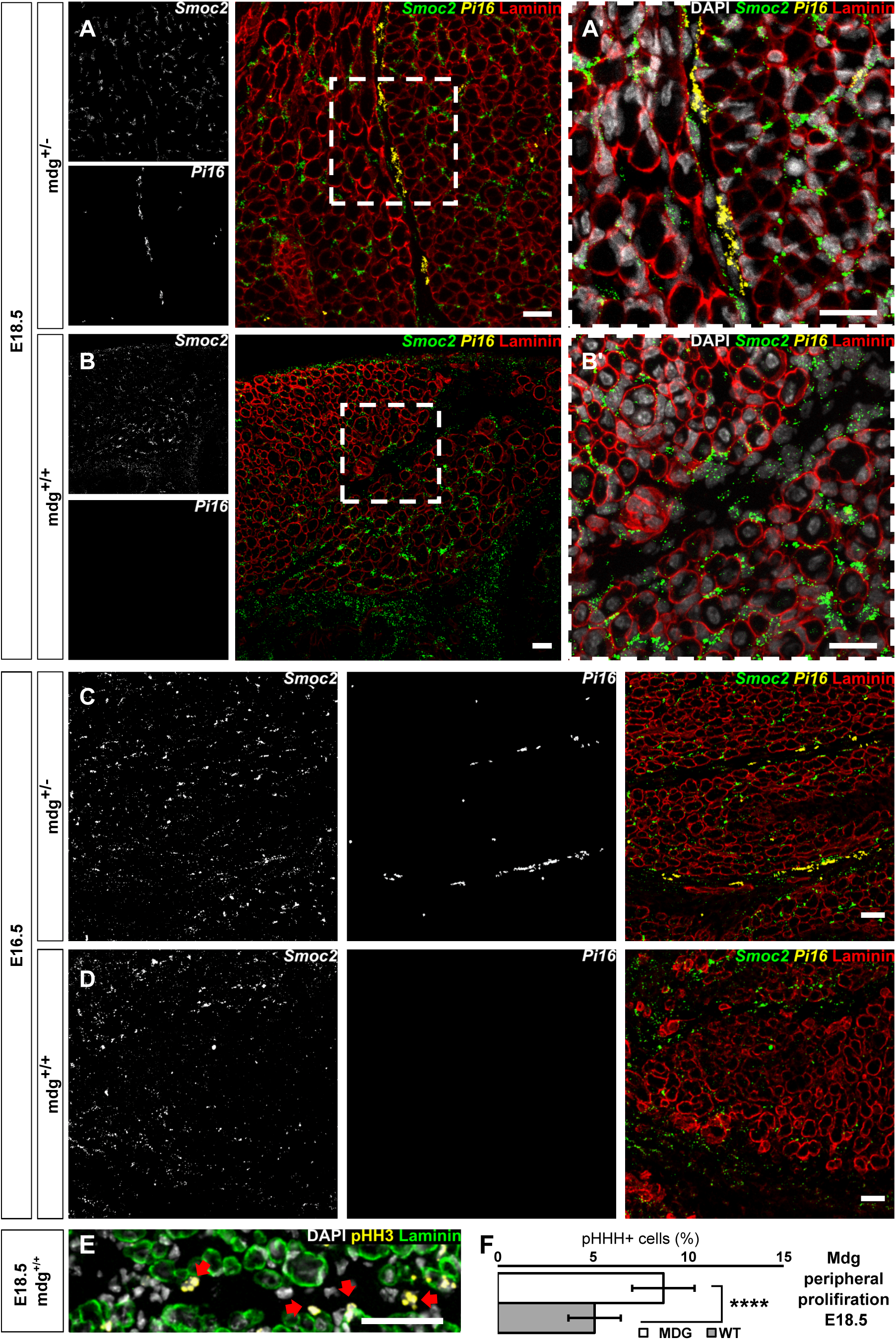
*Pi16+* fibroblasts are absent in muscular dysgenic mice. (A-D) Immunofluorescence staining and RNA FISH of fibroblast subpopulation markers (*Smoc2* and *Pi16*) in muscle cross sections across development at E18.5 (A-B) and E16.5 (C-D) in heterozygote (A,C) or homozygote (B,D) *mdg* embryos. DAPI (grey), Laminin (Red), *Smoc2* (green), *Pi16* (yellow). Scale bars = 25 µm. (E) Immunofluorescence staining for cell cycle marker pHH3 in muscle peripheral cells at E18.5 in homozygote *mdg* mutants. DAPI (gray), Laminin (green), pHH3 (yellow). Arrows highlight pHH3+ fibroblasts in muscle peripheral cells. Scale bars = 50 µm (F) Bar-plot quantification for pHH3+ muscle peripheral cells at E18.5 in *WT* and *mdg* mutants. n≥3 per group. Data are presented as mean ± SD. Significance denoted by an asterisk as calculated by two-tailed, unpaired Student’s t test.

To answer this question, we turned to an earlier developmental stage (E16.5), when *Pi16+/Smoc2+* dual expressing fibroblasts are abundant in muscle periphery (Fig. 4). At this stage, we found no expression of *Pi16* in the muscle periphery of *mdg* embryos (Fig. 6C,D). Furthermore, monitoring proliferation of peripheral fibroblasts at E18.5 in *mdg* muscles show an increased number of proliferating cells, supporting the idea of a disruption to the identity of these cells (Fig. 6E,F and Fig. S7F). Notably, Cldn1+ expression was maintained in perinerual fibroblasts in *mdg* mutant embryo (Fig. S7G). These data demonstrate that distinct cues regulate the diversification of distinct fibroblast subpopulations; while that of perineural fibroblasts is not dependent on muscle contraction, the initiation of the diversification of peripheral *Ly6a*+ from the *Smoc2*+ subpopulation relies on muscle contraction and does not occur in non-contractile muscles.

Overall, our results show that the development of tissue specific subpopulations seem to be independent from muscle contraction. In contrast, diversification of the ‘universal fibroblasts’ a developmental embryonic process, is dependent on muscle contraction, resulting in the formation of two large spatially segregated subpopulations which correspond to those found in adult muscles. These subpopulations occupy distinct niches in the limb, express unique transcriptomes including ECM and signaling related genes, and exert opposing effect on myogenic differentiation (Fig. 7).

**Figure 7.**
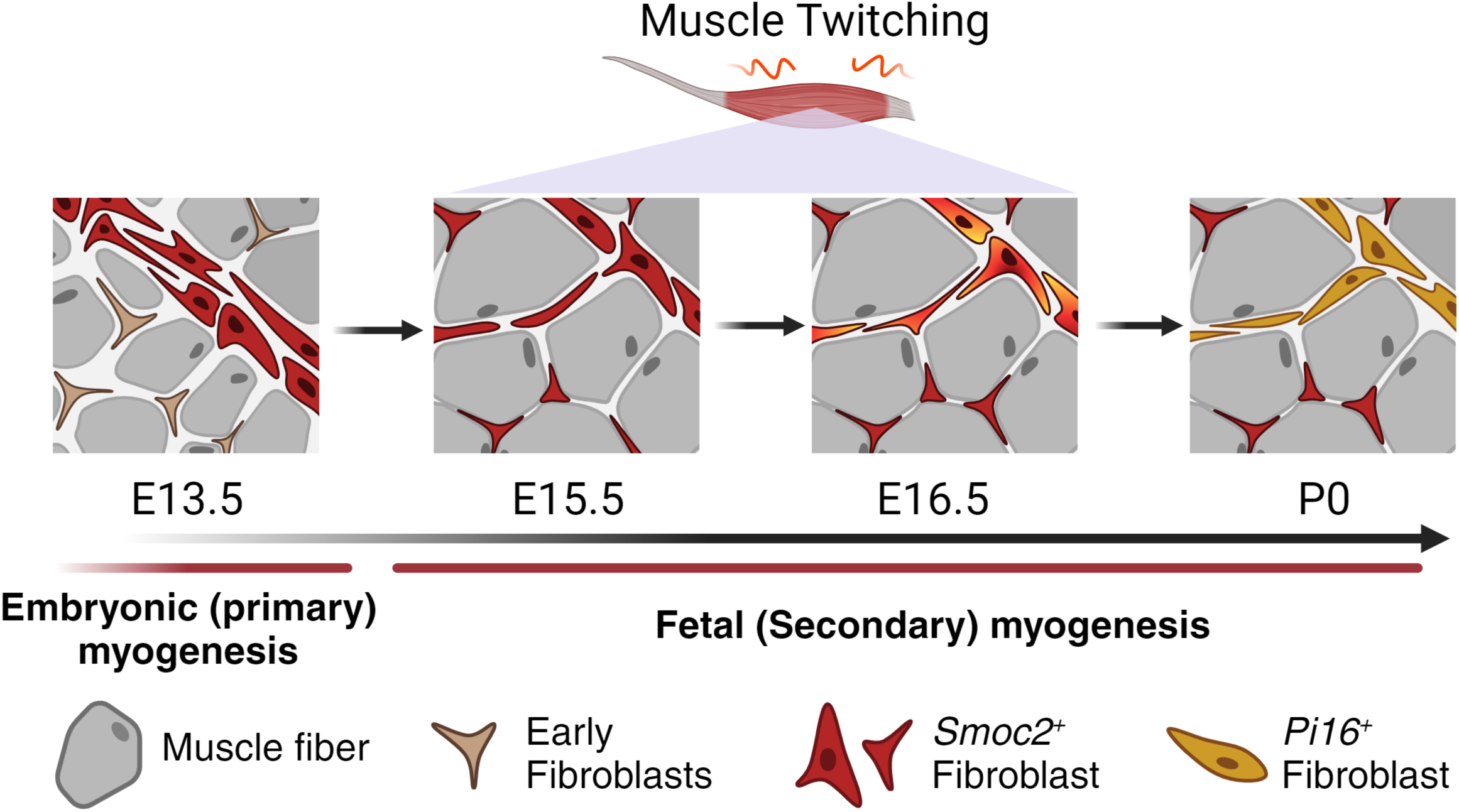
Model of embryonic diversification of fibroblasts. Working model of muscle resident fibroblast diversification during embryonic development, highlighting the role of muscle contractions in inducing the identity-transition of muscle delineating fibroblasts following the transition from embryonic to fetal myogenesis.

## DISSCUSSION

While it has become clear that fibroblasts are not a homogenous group of cells, the development and activities of the distinct subpopulations have not been characterized. Here, we focused on the two largest subpopulations, Pi16+ (Ly6a+) and Col15a1+ (Smoc2+), present in neonate muscles. We demonstrate fibroblast diversification is an embryonic process coinciding with the transition from primary to secondary myogenesis, segregating fibroblasts into spatially distinct subpopulations. This partition into interstitial Smoc2+ and delineating Ly6a+ fibroblasts is accompanied by transcriptional divergences, where each subpopulation acquires a distinct repertoire of differentially expressed marker genes and ECM proteins which correspond to their mature identities. Our work suggests these subpopulations differentially regulate myogenesis, where *Smoc2*+ fibroblasts promote myogenic proliferation while Ly6a+ fibroblasts inhibit differentiation. Our work further demonstrates that in conjunction with the transition from embryonic to fetal myogenesis, muscle contraction initiates the identity-transition of muscle delineating Smoc2+ fibroblasts into the Ly6a+ subtype. While in the muscle periphery, Ly6a+ fibroblasts halt their proliferation and deposit elastic ECM, interstitial Smoc2+ fibroblasts maintain higher proliferation rates, secrete components of muscle basement membrane, and promote myogenic proliferation to support myogenesis. We suggest this spatial and functional diversification facilitates the distinct roles of each fibroblast subtype in the embryo, which help direct and contribute to muscle development, possibly being carried on to the adult.

Embryonic myogenic progenitors or satellite cells in the adult, are muscle specific tissue resident stem cells capable of giving rise to skeletal muscle cells. These self-renewing cells undergo extensive proliferation in the embryo to facilitate developmental myogenesis, or in response to injury to muscle regeneration. In the muscle, these cells reside on top the myofibers and beneath the basal lamina in a specific ‘niche’, where they are presented with a plethora of signals and ques from the fiber or other cell types directing their fate. Extensive work has demonstrated fibroblasts to be essential for both embryonic myogenesis and adult regeneration, promoting myogenic proliferation and differentiation, as well as depositing and remodeling the muscle ECM (Contreras et al., 2021; Mathew et al., 2011; Molina et al., 2021; Nassari et al., 2017; Wosczyna et al., 2019). However, these works have approached fibroblasts as a uniform cell type, while recent studies dealing with fibroblast heterogeneity lack substantial functional data on subpopulation specific effects on myogenesis. Here, we are the first to show that fibroblast subpopulations exert distinct effects on myogenesis, as Smoc2+ fibroblasts promote the expansion of myogenic progenitors, while Ly6a+ fibroblasts inhibit myoblasts differentiation into mature myogenic cells. These findings support previous reports from transcriptomal studies suggesting Smoc2+ fibroblasts communicate with myogenic progenitors and support regeneration, supposedly by ECM remodeling (McKellar et al., 2021) although the observation that their effect on myogenic proliferation can be seen already following 24 hours of co-culture suggests other, ECM-independent mechanisms are involved. Altogether, when combined with our scRNAseq data and viewed through a spatial lens, our results suggest a new outlook at how these distinct subpopulations may differently contribute to regulating embryonic muscle development. Interstitial Smoc2+ fibroblasts found in close proximity to the myogenic progenitors produce basal membrane specific ECM proteins regulating signal presentation in the niche, and presumably secrete signaling molecules directly communicating with the myogenic progenitors (McKellar et al., 2021). Interestingly, although not evident in our data, few studies have reported the expression of retinoic acid related genes in interstitial fibroblasts (De Micheli et al., 2020a; Hirsinger et al., co-submitted; B. Zhang et al., 2023), suggesting a likely signaling pathway regulating the interactions between Smoc2 fibroblasts and myogenic progenitors (El Haddad et al., 2017; Lamarche et al., 2015; Zhao et al., 2020). In contrast, we found Ly6a+ fibroblasts to delineate the muscle and to highly express ECM proteins associated with elastic ECM fibers. Interestingly, Pi16 fibroblasts have been reported to reside in an adventitial layer and sheath other tissues beside muscle, making their delineating localization an inherent pan-tissue feature (Buechler et al., 2021). Thus, it could be hypothesized that these cells act as a size-limiting barrier, helping shape and instruct proper organ development.

FAPs are highly plastic multipotent mesenchymal progenitor cells which can give rise to both fibroblasts and adipogenic cells, closely linking the two lineages. Notably, recent work on the development of adipose tissue have identified a population of Dpp4+ adipose progenitors which delineate adipose depots. These Dpp4+ cells share many markers with Pi16+ fibroblasts (Jun et al., 2023; Merrick et al., 2019) suggesting they belong to the same subpopulation. However, no adipose related genes were significantly expressed in Pi16 fibroblasts in our data. Yet, the similar spatial distribution and transcriptomal signatures of Pi16 fibroblasts and Dpp4+ adipogenic progenitors, together with recent findings demonstrating Pi16 fibroblasts being highly adipogenic in vitro (Leinroth et al., 2022) highlights a possible role for Pi16 fibroblasts in postnatal adipogenesis and disease related muscle fatty infiltration. Interestingly, based on pseudotime analysis and the expression of stemness-related genes (Buechler et al., 2021; McKellar et al., 2021), *Pi16* fibroblasts have also been termed as ’stem’ FAPs in the adult, in keeping with our observation that these cells cease to proliferate following their introduction into their tissue-delineating niche.

Embryonic development can be described as a series of sequential processes dependent on prior and coordinated with parallel events, involving different tissues and cell types. In the early limb, a crosstalk between myogenic cells and the MCT orchestrates limb morphogenesis and patterning, promoting cell migration, expansion and differentiation in tightly timed processes (Helmbacher & Stricker, 2020). Our results indicate fibroblasts to have started diversifying by E15.5, accompanying the initiation of muscle contraction following the transition from primary to secondary myogenesis. Muscle contraction has been reported to be crucial for the development of several components of the musculoskeletal system in the embryo, including bone morphogenesis (Felsenthal & Zelzer, 2017), tendon patterning (Huang et al., 2015), MTJ and myofiber maturation (Dos Santos et al., 2023; Lipp et al., 2023). Results from the paralyzed *mdg* mutant showed loss of expression of Ly6a+ fibroblasts markers in the muscle periphery, suggesting that muscle contraction initiates these fibroblasts’ diversification. Our results further suggest that Cldn1+ fibroblasts develop independently from *Ly6a*+ fibroblasts, in agreement with differentiation trajectories demonstrating these subtypes diverge at different time-points (Arostegui et al., 2022). Additionally, analysis of transcriptional data from other models of paralyzed muscles (*RyR* and *Cacna1s* knockout) at E18.5 found a decrease in expression of genes associated with *Ly6a*+ fibroblasts, but not with *Smoc2+* or *Cldn1+* fibroblasts (Dos Santos et al., 2023; Filipova et al., 2018). These findings indicate that the failure to diversify is not inherent to the *mdg* model, supporting our hypothesis that *Ly6a*+ fibroblasts’ diversification is dependent on muscle contraction during development.

However, these results are not sufficient to determine whether muscle contraction directly triggers fibroblast diversification through mechanotransduction signaling, activated by biomechanical forces differently exerted on spatially distinct components of the limb, or indirectly, by promoting the maturation of other tissues instructing for fibroblast diversification. As we have discussed previously, Ly6a+ and Smoc2+ fibroblasts correspond to the ’universal subpopulations’ found in multiple tissues in our body (Buechler et al., 2021), some of which are not known to exert significant contractile forces as those observed in skeletal muscles. Therefore, an argument could be made for the role of secondary mediator signals driven by contractile forces, which induce fibroblasts diversification. Although the exact mechanism regulating this spatially restricted diversity awaits future work, deciphering the signals directing this process, in light of these subpopulations being highly evolutionary conserved from chick (Hirsinger et al., co-submitted) to humans (Buechler et al., 2021; De Micheli et al., 2020b; B. Zhang et al., 2023), could shed new light on the progression of fibrotic and connective tissue pathologies and serve as a basis for the development of new therapeutics combating the development of such ailments.

## Methods

### Mice

All experiments involving mice conform to the relevant regulatory standards (Technion IACUC and national animal welfare laws, guidelines and policies). Embryonic day was staged according to (Kaufman, 1992) ; noon on the day a vaginal plug was observed was marked as E0.5. For harvesting of embryos, timed-pregnant female mice were sacrificed by cervical dislocation.

For isolation of connective tissue cells, *Prrx1*^Cre^ line, targeting LPM derived cells (Logan & Tabin, 1998) were crossed with *Rosa26* ^nTnG^ as previously described (Yaseen et al., 2021). Mice heterozygous for the mutation muscular dysgenesis (mdg) (Pai, 1965) were obtained from the laboratory of Eli Zelzer, Rehovot, Israel. All mice are kept and bred on a C57Bl/6 background purchased from Envigo (https://www.envigo.com).

### Neonatal mouse muscle dissociation

P0 neonate limb muscles were sliced into pieces, incubated with 5 mg/ml collagenase II for 40 min at 37 °C in shaker, followed by mechanical dissociation using syringe. Tissues were then filtered, centrifuged for 51min at 300×g, re-suspended in PBS and counted.

### Fluorescent cell sorting

*WT* or *Prx1*^Cre^*; Rosa26*^nTnG^ P0 neonate limb muscles were dissociated and washed, cells were stained with antibodies staining mix for 30 min on ice, re-centrifuged and re-suspended in FACS sample medium (PBS + 2% 1% penicillin and streptomycin (PS)1+110% FCS + 2mM EDTA) and filtered before sorting.

Cells were then directly sorted using FACSAria™ IIIu (BD Biocsiences) and data analyzed using BD FACS Diva8.0.1 software. Gating strategy was the following: Cells were initially gated on FSC-A/SSC-A to exclude outliers and cell debris. Gated cells were then single gated twice based on FSC-W/FSC-H and then by SSC-W/SSC-H. For *Prx1*^Cre^*; Rosa26* ^nTnG^ muscles, single cells were then plotted on APC/Cy7 (CD45) versus APC (CD31) to remove immune and endothelial cells. Double negative cells were plotted on B530/30 (GFP) versus YG610/20 (RFP), and single GFP expressing cell were plotted on BV421 (CD200) versus PE/Cy7 (Sca1). Single RFP expressing cells (myogenic progenitors), single Sca1 expressing cells (*Ly6a*+ fibroblasts) and lineage negative cells (*Smoc2*+ fibroblasts) were sorted and. For *WT* muscles, single cells were then plotted on APC/Cy7 (CD45) versus APC (CD31 ; ITGA7) to remove immune, endothelial, and myogenic cells. Double negative cells were plotted on BV421 (CD200) versus PE/Cy7 (Sca1). Single Sca1 expressing cells (Ly6a+ fibroblasts) and lineage negative cells (Smoc2+ fibroblasts) were sorted and collected. Sorted cells were collected into BIO-AMF-2 (Biological Industries) + 2%PS

The following antibodies were used for sorting: anti-CD45 APC/Cy7 (103116; BioLegend), anti-CD31 APC (102410; BioLegend), anti-Sca1 PE/Cy7 (208114; BioLegend), anti-CD200 BV421 (565547; BD Horizon), and anti-ITGA7 APC (FAB3518A; R&D)

### Cell culture

Following sorting, fibroblasts were cultured on a plastic dish in fibroblast growth media (DMEM/F12[HAM];SATORIUS+10%FCS+ 1% L-glutamine and 1% PS). Myogenic cells cultured on Matrigel-coated plastic dish in BIO-AMF-2 (Biological Industries). Differentiation of myogenic cells was induced by incubating the cells in DMEM (containing 1% L-glutamine and 1% PS)1+14% horse serum.

### Immunofluorescence

10 µm fresh-frozen tissue sections were fixed using 4% paraformaldehyde, then blocked with 10% sheep serum or 1%BSA+1.5% donkey serum for anti-Pdgfra, and anti-Sca1 antibodies. Sections were incubated over-night at 4 °C or 2h at room temperature with 1st antibodies, followed by 2h incubation with 2nd antibodies.

Cell culture plates were fixed using 4% paraformaldehyde, and then blocked with 1% BSA.

The following antibodies were used in the immunostaining assays: Alexa Fluor® 488 Phalloidin (Cat# 8878, Cell Signaling, 1:200), anti-Myosin (Cat# A4.1025, DHSB, 1:300), anti-Laminin (Cat# L9393, Sigma, 1:400), anti-Collagen I (Cat# ab21286, Abcam, 1:200), anti-Sca1 (Cat# ab51317, Abcam, 1:100), anti-PDGFRα (Cat# AF1062, R&D, 1:100), anti-Cldn1 (Cat# ab115783, Abcam, 1:100), anti-Ly6c (Cat# ab54223, Abcam, 1:200), anti-CD31 (Cat# ab281583, Abcam, 1:100), anti-GFP (Cat# ab290, Abcam, 1:400).

All secondary antibodies were used at 1:200 dilution, nuclei were stained with DAPI.

### Immunohistochemistry

Whole-mount immunohistochemistry of neonate limbs was carried out as previously described in (Hasson et al., 2010), using anti-My32-AP (Cat# A4335, Sigma, 1:800)

### Fluorescent in situ hybridization (FISH)

FISH was carried out using the Molecular Instruments HCR-RNA-FISH system (Choi et al., 2018) according to the manufacturers’ protocol with slight modifications. Freshly cut frozen tissue sections were fixed using 4% paraformaldehyde for 10 min then incubated for 1 hour in 70% EtOH. Slides were acetylated using TEA and acetic anhydride, rinsed in DDW, and air dried for 30 min. After pre-hybridizing for 10 min, HCR probes (diluted 1:100) were added to the slides for overnight incubation at 37 °C. Slides were washed with Wash-Buffer 5xSSCT gradient then incubated with HCR amplifiers (diluted 1:50) overnight at room temperature. After the end of the protocol immunofluorescence was carried out as described previously.

### Imaging

Images were taken by Nikon Eclipse NI-E upright microscope or Zeiss LSM880 confocal microscope. Subsequently, all images were processed in a uniform way (average intensity projection, brightness adjustment and pseudo-coloring. RNA FISH images were passed through median filter for noise removal with radius = 0.2 pixels) using Fiji (Schindelin et al., 2012).

### 10× Genomics single cell general cell preparation

Single cell separation and library construction according to 10× protocol (Chromium Single Cell 31 Library & Gel Bead Kit v2). Briefly, skeletal muscles were dissected from E13.5 and 15.5 limbs, and single cells were isolated (as described) and suspended with BIO-AMF-2 (Biological Industries). The cells were mixed thoroughly using a wide-bore pipette tip and counted. Cells were centrifuged at 300×g for 5 min at RT, suspended with 0.04% BSA PBS, re-centrifuged and then re-suspended with the appropriate volume of 0.04% BSA PBS to achieve a target cell concentration in the range of 700–1200 cell/µl. Following cell isolation, library construction was immediately carried out using a Chromium Single Cell 3’ Reagent Kit (Version 3) and sequenced on an Illumina HiSeq 4000 or an Illumina NextSeq2000 (Illumina, cat no. 20040559) by the Technion Medicine Faculty Azrieli-Technion Genomics Center. Cell Ranger software (4.0.0) was used for data QC and extraction of transcripts’ counts from raw data (GEO accession <u>GSE259340</u>).

### Bioinformatic analysis

Seurat R package [Seurat 4.3.0,(Hao et al., 2021)] was used for filtering, clustering and expression distribution of selected cluster-specific genes of individual and integrated datasets. Cells with the following parameters were excluded: >8% mitochondrial UMI counts; less than 200 unique gene counts; over 4000 unique gene counts. Cells highly expressing Gm42418 and AY036118 were removed, as these genes overlap the rRNA element Rn45s and represent rRNA contamination, correlated with low sample quality (Akama-Garren et al., 2021) . Overall, 28,201 cells entered the analysis and 25,497 cells were used for the bioinformatics analysis after filtration. In addition, genes detected in less than three cells were filtered out. Integration of datasets into a single Seurat object were done using the Harmony R package [Harmony 0.1.0 (Korsunsky et al., 2019)]. For downstream clustering analysis, 30 PC and dimension number were chosen using seurat elbow and jackstraw plots as per Seurat’s standard workflow. Differential expression of relevant cell marker genes was visualized on UMAP plots to reveal specific individual cell types.

Cell cycle stage was predicted using Seurat’s CellCycleScoring function

Gene ontology (GO) and pathway analysis was performed using the online functional annotation tool g:Profiler (Raudvere et al., 2019). Collagen functional group scoring was following (Peisker et al., 2022) method. In short, functional group score was calculated using Seurat AddModuleScore function according to (Karsdal et al., 2020) functional classification of collagens.

Single-cell pseudotime trajectories were computed using the slingshot R package (version 2.4.0), which enables computation of lineage structures in a low-dimensional space (Street et al., 2018). Specifically, slingshot was implemented in the analysis pipeline after subsetting four clusters of interest of the fibroblast lineage, followed by dimensional reduction and clustering of the subsetted object. Pre-computed cell embeddings and clusters from the Seurat pipeline served as an input to the function. Mesenchymal progenitors were chosen as the start cluster using prior biological knowledge and the sample’s dating during pregnancy. For the analysis slingshot standard workflow was followed. Shortly, the wrapper function slingshot then performed lineage inference by treating clusters as nodes and constructing a minimum spanning tree (MST) between the nodes. Next, lineages or trajectories were defined by ordering clusters via tracing paths through the MST. Finally, individual pseudotime(s) were visualized using principal curves on the subseted fibroblast lineage.

### Data and code availability

Previously generated P0 scRNAseq data is available in the GEO repository under accession <u>GSE168153</u> (Yaseen et al., 2021). E12.5 and E15.5 scRNAseq data have been deposited at GEO under the accession number <u>GSE259340</u>.

No original code is reported for this manuscript.

### Graphics

All models and schematics were created with BioRender.com.

### Statistical Analyses

All data are represented as mean ± SD. Statistical analyses were carried out using Microsoft Exel software or *Statistics Kingdom 2017 web application* . Unpaired two tailed t-tests and One-way anova were used as stated in the figure legends. For both in-vivo and in-vitro analyses, minimal number of 3 mice or experimental repeats were used per group, listed in figure legends as n. For all cell isolation experiments (FACS and scRNA sequencing), a pool of cells from at least 4 embryos were mixed for sorting/sequencing. Significance was considered with a minimum p value of 0.05, and denoted by a number of asterisks: p < 0.05 = *, p < 0.01 = **, p < 0.001 = ***, p < 0.0001 = ****.

For myogenic differentiation, Fusion index was quantified as previously described (total number of nuclei within myofibers with IZ3 nuclei/total number of nuclei within all myofibers) ×100 (Mathew et al., 2011).

## Author Contribution

L.C. conducted the majority of the experiments. S.Z.-E. assisted in designing and conducting cell isolation and culture experiments. A.K. assisted in cell culture experiments and immunofluorescence staining. A.O. participated in image processing and analysis. L.C and P.H supervised and analyzed the data. L.C. and P.H. planned the study and wrote the manuscript.

## Acknowledgments

We are grateful to M. Holdengreber and M. Gurewitz from the Biomedical core facilities (BCF) Light Microscopy unit (Faculty of Medicine, Technion); to A. Grau and J. Bisharat from the BCF Cytometry unit (Faculty of Medicine, Technion); to L. Linde and M. Peleg from the Azrieli-Technion Genomics Center (Faculty of Medicine, Technion) for bioinformatics analyses; to members of the animal facility for excellent technical assistance. P.H. was supported by grants from the Israeli Science Foundation (1111/18 and 1126/23), by A.D.I. (Association Duchenne Israel) and by the Rappaport Family Institute.

## Notes

### Competing Interest Statement

The authors have declared no competing interest.

