## supplemental Figures for "Fibroblast diversification is an embryonic process dependent on muscle contraction"

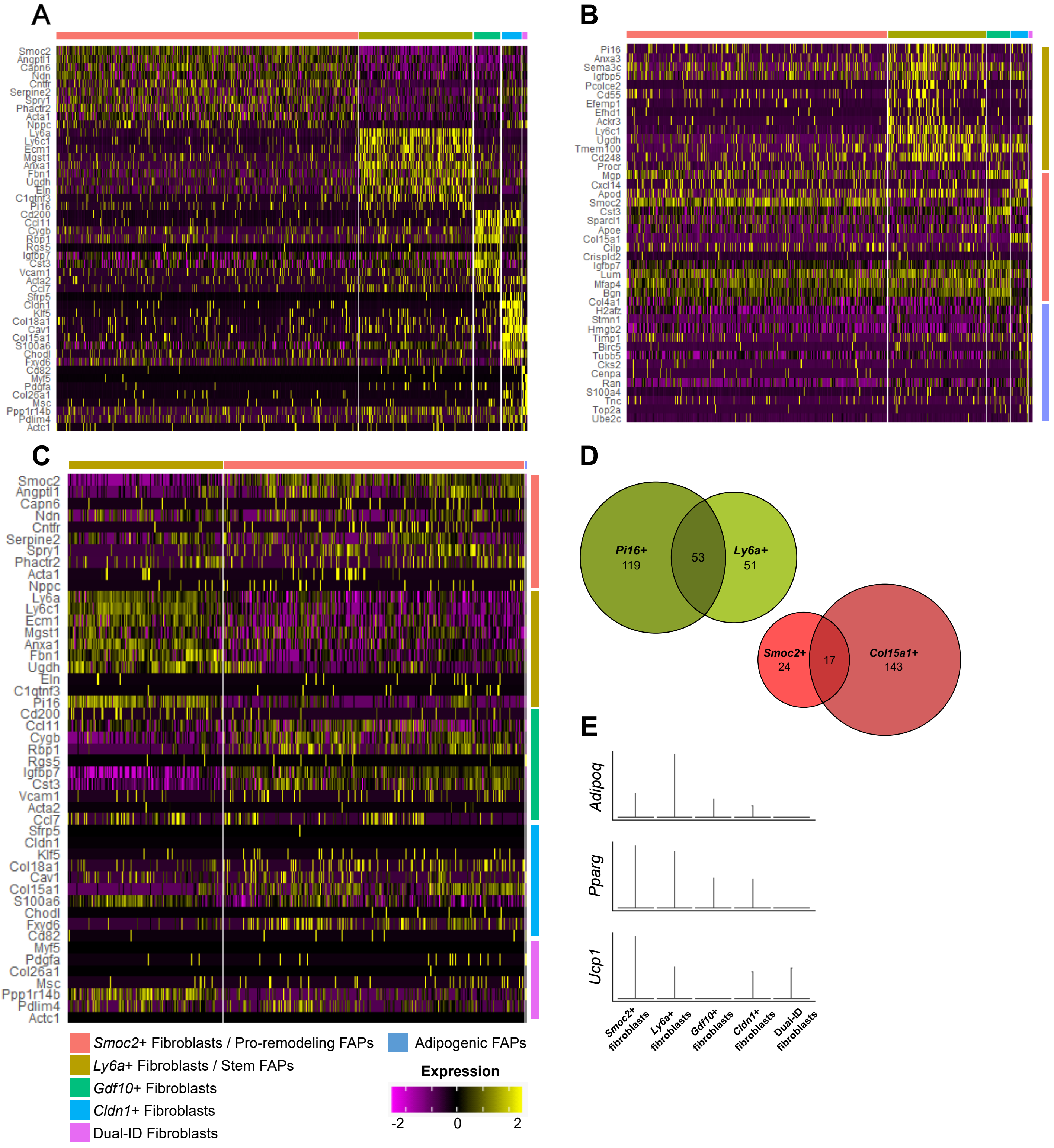
A
Smoc2
Angptl1
Capn6
Ndn
Cntfr
Serpine2
Spry1
Phactr2
Acta1
Nppc
Ly6a
Ly6c1
Ecm1
Mgst1
Anxa1
Fbn1
Ugdh
Eln
C1qtnf3
Pi16
Cd200
Ccl11
Cygb
Rbp1
Rgs5
Igfbp7
Cst3
Vcam1
Acta2
Ccl7
Sfrp5
Cldn1
Klf5
Col18a1
Cav1
Col15a1
S100a6
Chodl
Fxyd6
Cd82
Myf5
Pdgfa
Col26a1
Msc
Ppp1r14b
Pdlim4
Actc1
B
Pi16
Anxa3
Sema3c
Igfbp5
Pcolce2
Cd55
Efemp1
Efhd1
Ackr3
Ly6c1
Ugdh
Tmem100
Cd248
Procr
Mgp
Cxcl14
Apod
Smoc2
Cst3
Sparcl1
Apoe
Col15a1
Cilp
Crispld2
Igfbp7
Lum
Mfap4
Bgn
Col4a1
H2afz
Stmn1
Hmgb2
Timp1
Birc5
Tubb5
Cks2
Cenpa
Ran
S100a4
Tnc
Top2a
Ube2c
C
Smoc2
Angptl1
Capn6
Ndn
Cntfr
Serpine2
Spry1
Phactr2
Acta1
Nppc
Ly6a
Ly6c1
Ecm1
Mgst1
Anxa1
Fbn1
Ugdh
Eln
C1qtnf3
Pi16
Cd200
Ccl11
Cygb
Rbp1
Rgs5
Igfbp7
Cst3
Vcam1
Acta2
Ccl7
Sfrp5
Cldn1
Klf5
Col18a1
Cav1
Col15a1
S100a6
Chodl
Fxyd6
Cd82
Myf5
Pdgfa
Col26a1
Msc
Ppp1r14b
Pdlim4
Actc1
D
Pi16+
119
53
Ly6a+
51
Smoc2+
24
17
Col15a1+
143
E
Adipoq
Pparg
Ucp1
Smoc2+ fibroblasts
Ly6a+ fibroblasts
Gdf10+ fibroblasts
Cldn1+ fibroblasts
Dual-ID fibroblasts
Smoc2+ Fibroblasts / Pro-remodeling FAPs
Ly6a+ Fibroblasts / Stem FAPs
Gdf10+ Fibroblasts
Cldn1+ Fibroblasts
Dual-ID Fibroblasts
Adipogenic FAPs
Expression
-2
0
2

**Figure S1. Neonate fibroblast clusters correspond to adult populations**

(A) Heatmap of scRNA-seq data showing the top 10 enriched genes in the 5 different clusters at P0. Identity of expressing cluster is noted along the top (with corresponding cluster identity at the bottom).

(B) Heatmap of scRNA-seq data showing expression of the top 15 DEGs of adult fibroblast clusters in the 5 P0 fibroblast clusters.

(C) Heatmap of scRNA-seq data showing expression of the top 10 DEGs of P0 fibroblasts in the 3 different non-injured adult fibroblasts clusters.

(B-C) Identity of expressing cluster is noted along the top and identity of the gene-sets' cluster origin is noted along the right (with corresponding cluster identity at the bottom).

(D) Venn diagram depicting highly DEGs ($log_2FC \geq 1$) comparisons between P0 Smoc2+ and Ly6a+ fibroblast clusters with non-injured adult Pi16+ and Col15a1+ fibroblast clusters.

(E) Violin plots for Adipogenic hallmark genes in the 5 fibroblast clusters at P0.

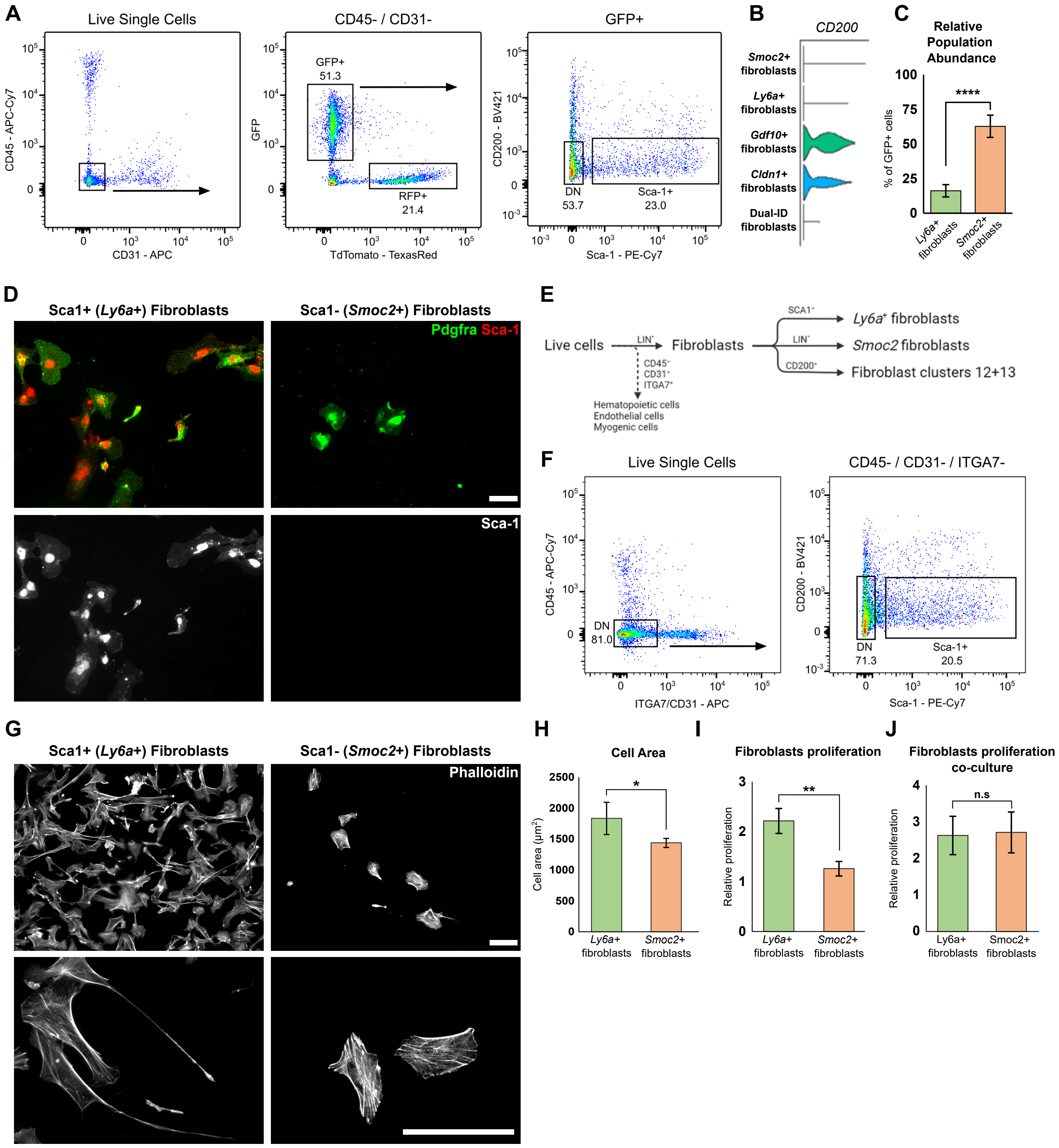
A
Live Single Cells
CD45 - APC-Cy7
CD31 - APC
CD45- / CD31-
GFP+
51.3
RFP+
21.4
GFP
TdTomato - TexasRed
GFP+
CD200 - BV421
DN
53.7
Sca-1+
23.0
Sca-1 - PE-Cy7
B
CD200
Smoc2+ fibroblasts
Ly6a+ fibroblasts
Gdf10+ fibroblasts
Cldn1+ fibroblasts
Dual-ID fibroblasts
C
Relative Population Abundance
% of GFP+ cells
****
Ly6a+ fibroblasts
Smoc2+ fibroblasts
D
Sca1+ (Ly6a+) Fibroblasts
Sca1- (Smoc2+) Fibroblasts
Pdgfra Sca-1
Sca-1
E
Live cells
LIN-
Fibroblasts
SCA1+
Ly6a+ fibroblasts
LIN-
Smoc2 fibroblasts
CD200+
Fibroblast clusters 12+13
CD45+
CD31+
ITGA7+
Hematopoietic cells
Endothelial cells
Myogenic cells
F
Live Single Cells
CD45 - APC-Cy7
DN
81.0
ITGA7/CD31 - APC
CD45- / CD31- / ITGA7-
CD200 - BV421
DN
71.3
Sca-1+
20.5
Sca-1 - PE-Cy7
G
Sca1+ (Ly6a+) Fibroblasts
Sca1- (Smoc2+) Fibroblasts
Phalloidin
H
Cell Area
Cell area (µm²)
*
Ly6a+ fibroblasts
Smoc2+ fibroblasts
I
Fibroblasts proliferation
Relative proliferation
**
Ly6a+ fibroblasts
Smoc2+ fibroblasts
J
Fibroblasts proliferation co-culture
Relative proliferation
n.s
Ly6a+ fibroblasts
Smoc2+ fibroblasts

**Figure S2. Smoc2+ and Ly6a+ fibroblast subpopulations show distinct properties in vitro**

(A) FACS plots showing isolation of cell populations from *$Prx1^{Cre}$;$ROSA^{nTnG}$* P0 limb muscle cells. DN = double negative.

(B) Violin plot showing expression of CD200 in fibroblast subpopulations at P0.

(C) Percent quantification of relative fibroblast subpopulation abundance from connective tissue lineage (GFP+ cells).

(D) Immunostaining of sorted Ly6a+ and Smoc2+ fibroblasts for general fibroblastic (Pdgfra) and specific (Sca-1) markers. Pdgfra (green), Sca-1 (red). Scale bar = 100 µm.

(E) Flowchart schematic showing sorting fibroblasts sorting strategy from WT postnatal limb muscles.

(F) FACS plots showing isolation of fibroblast subpopulations from WT postnatal limb muscle cells. DN = double negative.

(G) Phalloidin staining of sorted Ly6a+ and Smoc2+ after 24h in culture at low (10X, upper) and high (40X, bottom) magnifications. Phalloidin (gray). Scale bar = 100 µm.

(H) Mean cell area of sorted cells area after 24 hours. n = 3.

(I) Proliferation rates of fibroblast subpopulations over 60 hour culturing, relative to initial density of seeded cells

(J) Proliferation rates of fibroblast subpopulations over 96 hour co-culturing with myogenic progenitors, relative to initial density of seeded cells.

(C, H-J) Data are presented as mean ± SD. Significance denoted by an asterisk as calculated by two-tailed, unpaired Student's t test.

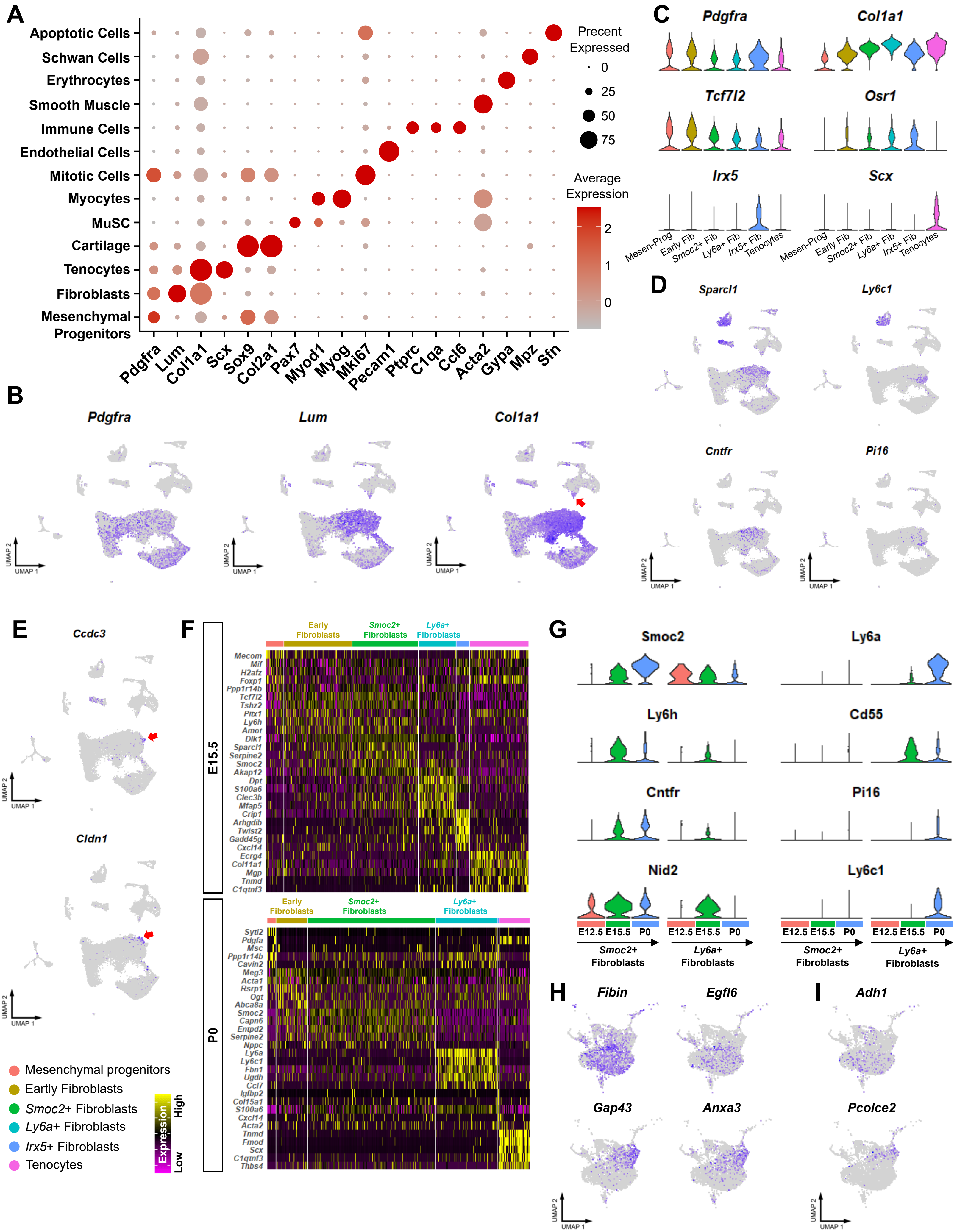
A
Apoptotic Cells
Schwan Cells
Erythrocytes
Smooth Muscle
Immune Cells
Endothelial Cells
Mitotic Cells
Myocytes
MuSC
Cartilage
Tenocytes
Fibroblasts
Mesenchymal Progenitors
Pdgfra Lum Col1a1 Scx Sox9 Col2a1 Pax7 Myod1 Myog Mki67 Pecam1 Ptprc C1qa Ccl6 Acta2 Gypa Mpz Sfn
Precent Expressed
0
25
50
75
Average Expression
2
1
0
B
Pdgfra
Lum
Col1a1
UMAP 2
UMAP 1
C
Pdgfra
Col1a1
Tcf7l2
Osr1
Irx5
Scx
Mesen-Prog
Early Fib
Smoc2+ Fib
Ly6a+ Fib
Irx5+ Fib
Tenocytes
D
Sparcl1
Ly6c1
Cntfr
Pi16
E
Ccdc3
Cldn1
F
Early Fibroblasts
Smoc2+ Fibroblasts
Ly6a+ Fibroblasts
E15.5
Mecom
Mif
H2afz
Foxp1
Ppp1r14b
Tcf7l2
Tshz2
Pitx1
Ly6h
Amot
Dlk1
Sparcl1
Serpine2
Smoc2
Akap12
Dpt
S100a6
Clec3b
Mfap5
Crip1
Arhgdib
Twist2
Gadd45g
Cxcl14
Ecrg4
Col11a1
Mgp
Tnmd
C1qtnf3
P0
Sytl2
Pdgfa
Msc
Ppp1r14b
Cavin2
Meg3
Acta1
Rsrp1
Ogt
Abca8a
Smoc2
Capn6
Entpd2
Serpine2
Nppc
Ly6a
Ly6c1
Fbn1
Ugdh
Ccl7
Igfbp2
Col15a1
S100a6
Cxcl14
Acta2
Tnmd
Fmod
Scx
C1qtnf3
Thbs4
Mesenchymal progenitors
Eartly Fibroblasts
Smoc2+ Fibroblasts
Ly6a+ Fibroblasts
Irx5+ Fibroblasts
Tenocytes
Expression
High
Low
G
Smoc2
Ly6a
Ly6h
Cd55
Cntfr
Pi16
Nid2
Ly6c1
E12.5 E15.5 P0
Smoc2+ Fibroblasts
Ly6a+ Fibroblasts
H
Fibin
Egfl6
Gap43
Anxa3
I
Adh1
Pcolce2

**Figure S3. Temporal deconstruction of fibroblast heterogeneity and diversification during development**

(A) Dot plot of canonical markers of cell types showing cluster-average expression. Dot size represents the percentage of cells with a non-zero expression level and color-scale represents the average expression level across all cells within cluster.

(B) UMAPs of canonical fibroblastic markers in integrated dataset. Red arrow highlights Dual-ID fibroblasts (Col1a1 expressing cells clustering with MuSC).

(C) Violin plots for general fibroblast markers and cluster specific markers (bottom row) in connective tissue lineage subset from integrated dataset. Mesenchymal Progenitors (Mesen-Prog).

(D) UMAPs of characteristic markers of *Smoc2*+ (left) and *Ly6a+* (right) fibroblast subpopulations in integrated dataset.

(E) UMAPs of characteristic markers of *Gdf10*+ (top) and *Cldn1* (bottom) fibroblast subpopulations in integrated dataset. Red arrow highlight expressing cells.

(F) Heatmap of scRNA-seq data showing the top 10 enriched genes in different connective tissue lineage clusters at E15.5 (top) and P0 (bottom) . Identity of expressing cluster is noted along the top (with corresponding cluster identity at the bottom).

(G) Violin plots for expression of Smoc2+ and Ly6a+ marker genes in *Smoc2*+ (left), and *Ly6a*+ (right) fibroblasts in each timepoint. Colors represent developmental stage of origin as in (Fig. 3C).

(H, I) UMAPs of characteristic markers of fibroblast subpopulations corresponding to *Smoc2*+ (top) and *Ly6a* (bottom) fibroblasts identified in chick (H) and human (I).

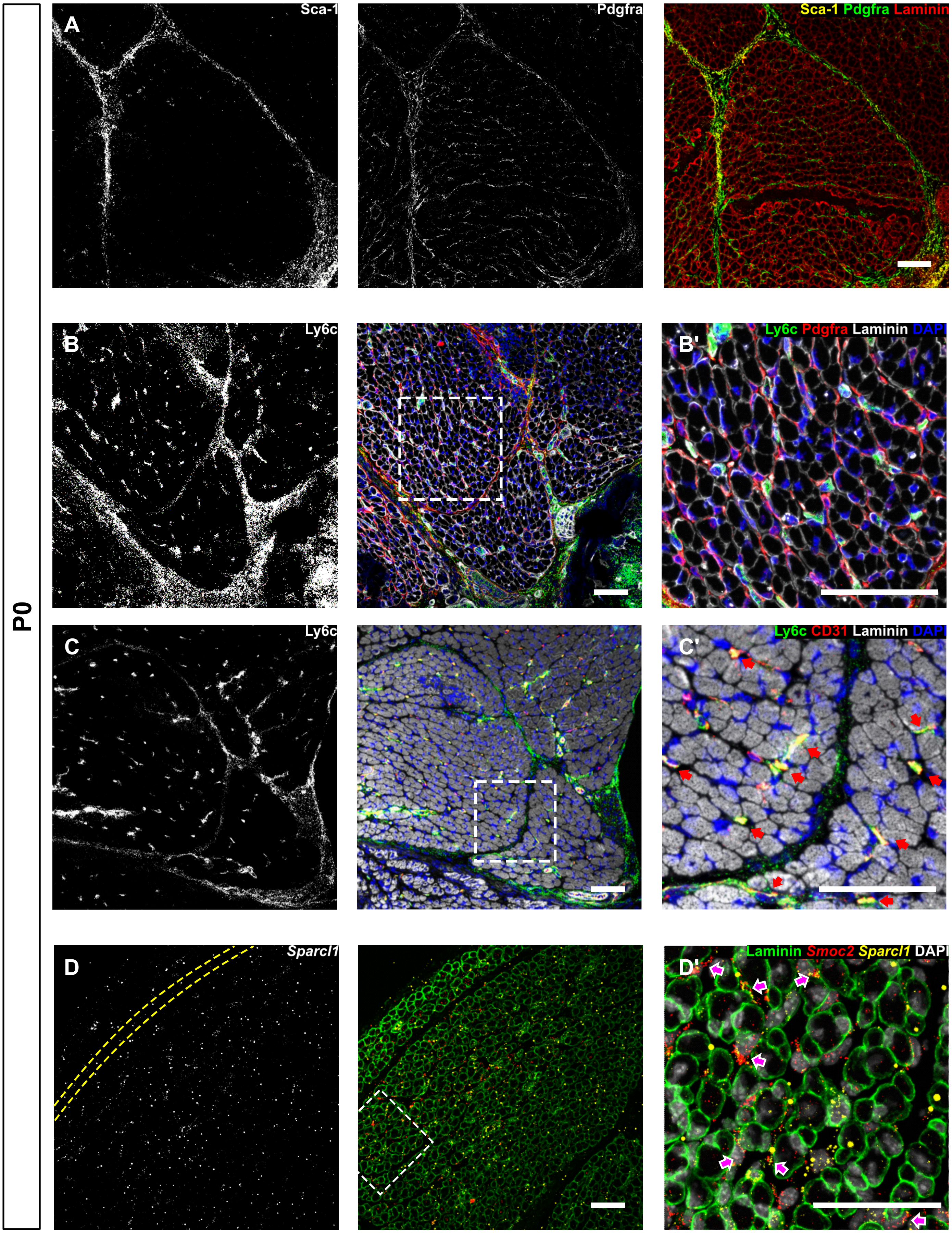

P0
A
Sca-1
Pdgfra
Sca-1 Pdgfra Laminin
B
Ly6c
B'
Ly6c Pdgfra Laminin DAPI
C
Ly6c
C'
Ly6c CD31 Laminin DAPI
D
Sparcl1
D'
Laminin Smoc2 Sparcl1 DAPI

**Figure S4. Spatial diversification of fibroblast heterogeneity during development**

(A-C') Immunofluorescence staining of *Ly6a*+ fibroblast subpopulation markers in muscle cross sections at P0. Arrows highlight CD31+ and Ly6c+ blood vessels (C').

(D-D') Immunofluorescence staining and RNA FISH of *Smoc2*+ fibroblast subpopulation markers in muscle cross sections at P0. The muscle borders are highlighted by a yellow dashed line (D). Arrows highlight *Smoc2*+ and *Sparcl1*+ interstitial fibroblasts (D').

Scale bars = 50 µm.

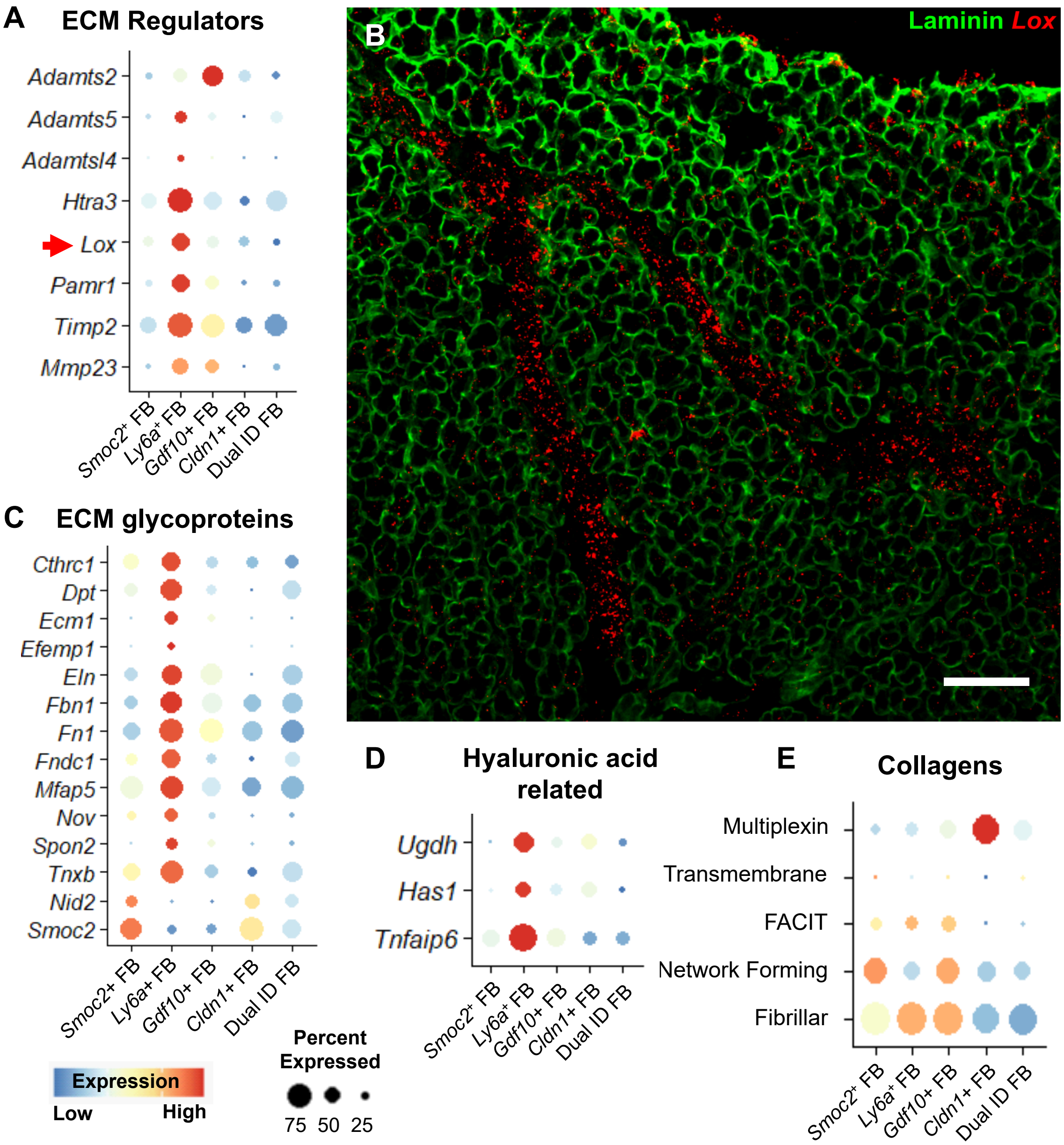
A
ECM Regulators
Adamts2
Adamts5
Adamtsl4
Htra3
Lox
Pamr1
Timp2
Mmp23
Smoc2+ FB
Ly6a+ FB
Gdf10+ FB
Cldn1+ FB
Dual ID FB
B
Laminin Lox
C
ECM glycoproteins
Cthrc1
Dpt
Ecm1
Efemp1
Eln
Fbn1
Fn1
Fndc1
Mfap5
Nov
Spon2
Tnxb
Nid2
Smoc2
Smoc2+ FB
Ly6a+ FB
Gdf10+ FB
Cldn1+ FB
Dual ID FB
Expression
Low
High
Percent
Expressed
75
50
25
D
Hyaluronic acid
related
Ugdh
Has1
Tnfaip6
Smoc2+ FB
Ly6a+ FB
Gdf10+ FB
Cldn1+ FB
Dual ID FB
E
Collagens
Multiplexin
Transmembrane
FACIT
Network Forming
Fibrillar
Smoc2+ FB
Ly6a+ FB
Gdf10+ FB
Cldn1+ FB
Dual ID FB

**Figure S5. Expression of ECM and ECM related genes in fibroblasts subpopulations in neonate muscle**

(A) Dot plot showing ECM regulators gene expression in fibroblast clusters at P0. Dot size represents the percentage of cells with a non-zero expression level and color-scale represents the average expression level across all cells within the cluster.

(B) Immunofluorescence staining and RNA FISH of ECM regulator *Lox* gene in muscle cross sections at E18.5. Laminin (green), *Lox* (red). Scale bar = 50 µm

(C-E) Dot plot showing expression of ECM glycoproteins (C), Hyaluronic acid related (D) genes and collagen functional groups (E) in fibroblast clusters at P0. Dot size represents the percentage of cells with a non-zero expression level and color-scale represents the average expression level across all cells within cluster.

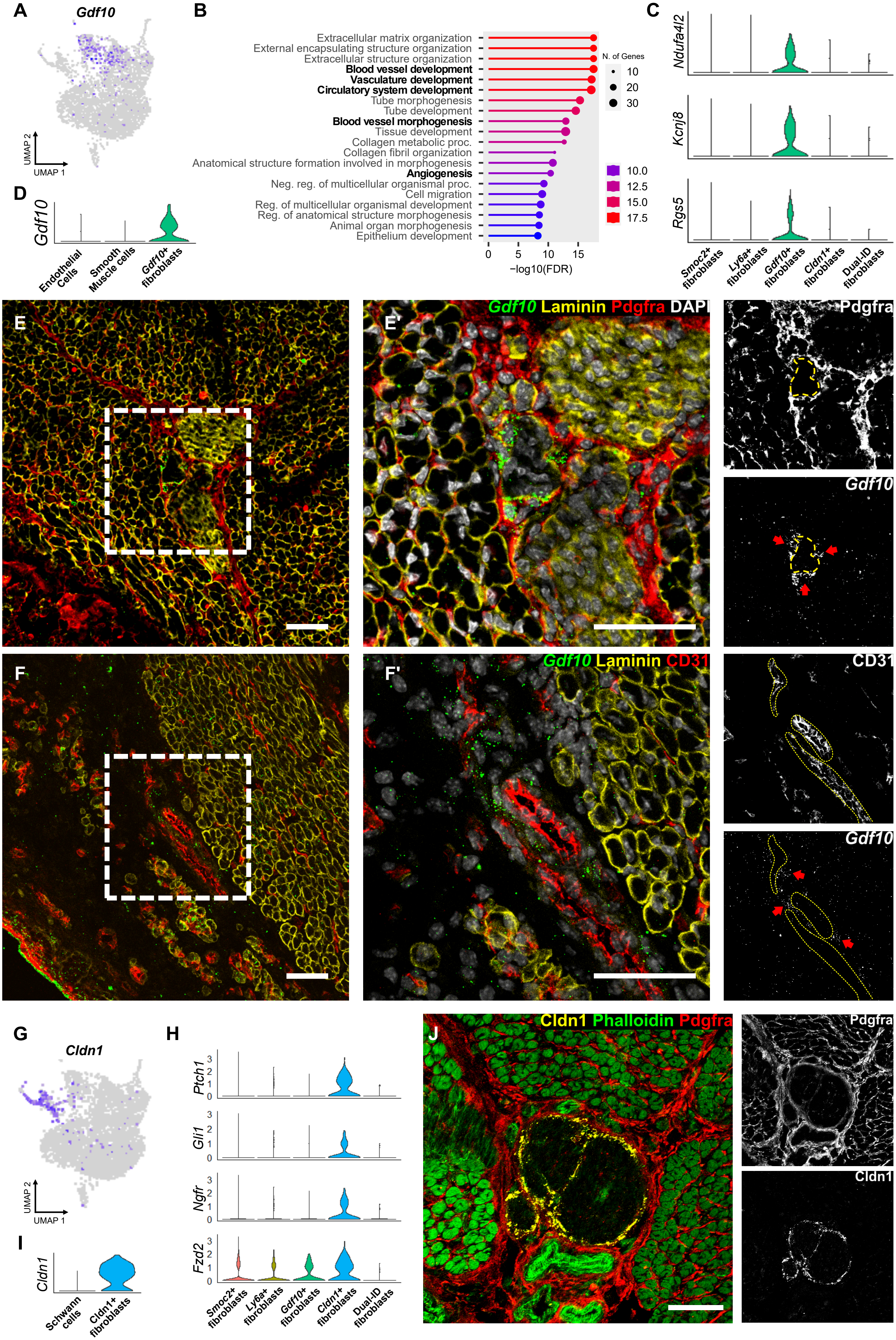

A
Gdf10
UMAP 2
UMAP 1
B
Extracellular matrix organization
External encapsulating structure organization
Extracellular structure organization
Blood vessel development
Vasculature development
Circulatory system development
Tube morphogenesis
Tube development
Blood vessel morphogenesis
Tissue development
Collagen metabolic proc.
Collagen fibril organization
Anatomical structure formation involved in morphogenesis
Angiogenesis
Neg. reg. of multicellular organismal proc.
Cell migration
Reg. of multicellular organismal development
Reg. of anatomical structure morphogenesis
Animal organ morphogenesis
Epithelium development
0 5 10 15
−log10(FDR)
N. of Genes
10
20
30
10.0
12.5
15.0
17.5
C
Ndufa4l2
Kcnj8
Rgs5
Smoc2+ fibroblasts
Ly6a+ fibroblasts
Gdf10+ fibroblasts
Cldn1+ fibroblasts
Dual-ID fibroblasts
D
Gdf10
Endothelial Cells
Smooth Muscle cells
Gdf10+ fibroblasts
E
E'
Gdf10 Laminin Pdgfra DAPI
Pdgfra
Gdf10
F
F'
Gdf10 Laminin CD31
CD31
Gdf10
G
Cldn1
UMAP 2
UMAP 1
H
Ptch1
Gli1
Ngfr
Fzd2
0 1 2 3
Smoc2+ fibroblasts
Ly6a+ fibroblasts
Gdf10+ fibroblasts
Cldn1+ fibroblasts
Dual-ID fibroblasts
I
Cldn1
Schwann cells
Cldn1+ fibroblasts
J
Cldn1 Phalloidin Pdgfra
Pdgfra
Cldn1

**Figure S6. Characterization and spatial localization of *Gdf10*+ and Cldn1+ fibroblast subpopulations**

(A) UMAP gene expression for *Gdf10* demonstrate specific expression of the marker in distinct fibroblast clusters at P0.

(B) Go term analysis of the top 100 DEGs in Gdf10+ fibroblasts. Vascular system related terms are in bold. Dot size represents the number of genes identified per term. Color-scale represents the –log10(FDR) of each term.

(C) Violin plots showing specific expression of pericyte marker genes in *Gdf10*+ fibroblasts at P0.

(D) Violin plot for *Gdf10* demonstrate specific expression in fibroblasts and not other vascular cells, namely endothelial and smooth muscle, at P0.

(E-F') Immunofluorescence staining and RNA FISH of *Gdf10* gene in blood vessel associated fibroblasts in muscle cross sections at P0. An orange dashed line highlights blood vessel borders. Arrows highlight *Gdf10* expressing fibroblasts adjacent to blood vessels. Laminin (yellow), *Gdf10* (green), DAPI (white), Pdgfra (E-E') or CD31 (F-F') (red). Scale bars = 50 µm.

(G) UMAP gene expression for *Cldn1* demonstrate specific expression of the marker in a distinct fibroblast cluster at P0.

(H) Violin plots showing specific expression of *Cldn1* perineural fibroblast marker genes in *Cldn1*+ fibroblasts at P0.

(I) Violin plot for *Cldn1* demonstrate specific expression of the marker in nerve related fibroblasts but not in Schwan cells at P0.

(J) Immunofluorescence staining of Cldn1 and Pdgfrα in perineural fibroblasts in muscle cross sections at P0. Phalloidin (green), Cldn1 (yellow), Pdgfrα (red). Scale bars = 50 µm.

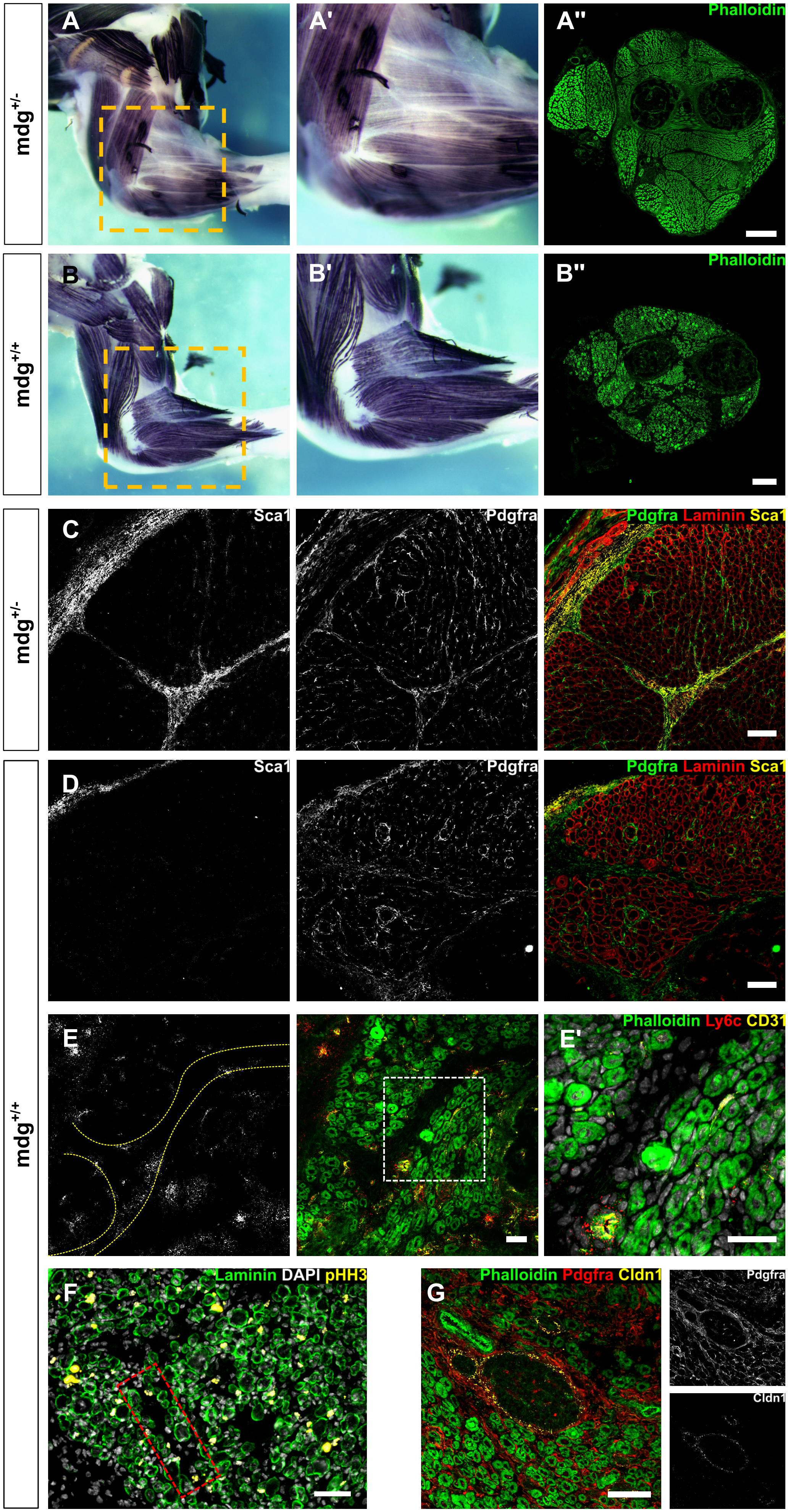
mdg+/-
A
A'
A''
Phalloidin
mdg+/+
B
B'
B''
Phalloidin
mdg+/-
C
Sca1
Pdgfra
Pdgfra Laminin Sca1
mdg+/+
D
Sca1
Pdgfra
Pdgfra Laminin Sca1
E
E'
Phalloidin Ly6c CD31
F
Laminin DAPI pHH3
G
Phalloidin Pdgfra Cldn1
Pdgfra
Cldn1

**Figure S7. Impairment of muscle maturation and fibroblast subtype marker expression in paralyzed muscles.**

(A-B’’) Wholemount immunostaining for myosin heavy chain (A-A’,B-B’) and phalloidin (green) section fluorescent staining (A’’,B’’) of heterozygote (A-A’’) or homozygote (B-B’’) *mdg* mutant embryo limbs at E18.5. Scale bars = 200 µm.

(C-E’) Immunofluorescence staining for Sca1 marking *Ly6a*+ fibroblast subpopulation in heterozygote (C) and homozygote (D-E’) *mdg* muscle cross sections at E18.5. Muscle borders are highlighted by a yellow dashed line (E). Pdgfra (green), Laminin (red) and Sca1 (yellow) (C-D). Phalloidin (green), Ly6c (red) and CD31 (yellow) (E-E’) DAPI (E’). Scale bars = 50 µm.

(F) Lower magnification for Fig. 6E. Immunofluorescence staining for cell cycle marker pHH3 in muscle peripheral cells at E18.5 in homozygote *mdg* mutants. DAPI (gray), Laminin (green), pHH3 (yellow). Scale bar = 50 µm.

(G) Immunofluorescence staining of Cldn1 and Pdgfrα in perineural fibroblasts in mdg muscle cross sections at E18.5. Phalloidin (green), Cldn1 (yellow), Pdgfrα (red). Scale bars = 50 µm.
